# Structural Genomics of the Human Dopamine Receptor System

**DOI:** 10.1101/2022.10.09.511478

**Authors:** Peiyu Xu, Sijie Huang, Brian E. Krumm, Youwen Zhuang, Chunyou Mao, Yumu Zhang, Yue Wang, Xi-Ping Huang, Yong-Feng Liu, Xinheng He, Huadong Li, Wanchao Yin, Yi Jiang, Yan Zhang, Bryan L. Roth, H. Eric Xu

## Abstract

The dopamine system, including five dopamine receptors (D1R to D5R), plays essential roles in the central nervous system (CNS) and ligands that activate dopamine receptors have been used to treat many neuropsychiatric disorders, including Parkinson’s Disease (PD) and schizophrenia. Here, we report five cryo-EM structures of all subtypes of human dopamine receptors in complex with G-protein and bound to the pan agonist, Rotigotine, which is used to treat PD and restless legs syndrome. The structures reveal the basis of Rotigotine binding modes to different dopamine receptors. Structural analysis together with functional assays illuminate determinants of ligand polypharmacology and selectivity. The structures also uncover the mechanisms of the dopamine receptor activation, unique structural features among the five receptor subtypes, and the basis of G-protein coupling specificity. Our works provide a comprehensive set of structural templates for the rational design of specific ligands to treat CNS diseases targeting the dopaminergic system.

## INTRODUCTION

Dopamine and the dopamine receptor system play critical roles in motor functions, cognition, and addiction ^1–3^. The action of dopaminergic system is mediated by five subtypes of dopamine receptors, a subfamily of G-protein coupled receptors (GPCRs). The dopamine receptors are divided into D1-like and D2-like groups. The D1-like group includes D1R and D5R, whereas the D2-like group includes D2R, D3R, and D4R. D1-like receptors are coupled to the stimulatory G-proteins (G_s_) and linked to the activation of adenylate cyclase. The D2-like receptors are coupled to the inhibitory subtypes of G-proteins (G_i_ and G_o_) and linked to the inhibition of adenylate cyclase ^4^.

Dopamine receptors are a prototypical class of drug targets for many CNS diseases, including Parkinson’s disease ^5^, schizophrenia ^6^, and attention deficit hyperactivity disorder (ADHD) ^7^. There are several dozens of dopaminergic drugs ^8^, with many of them having distinct properties of polypharmacology, which can act on multiple dopamine receptors or even other types of monoamine neurotransmitter receptors. However, understanding the polypharmacology of dopaminergic drugs remains a tremendous challenge due to the promiscuous binding of drugs to many different receptors with various pharmacology. Rotigotine, a drug for PD and restless legs syndrome (RLS), is a pan agonist that activates all five dopamine receptors ^9,10^. The molecular basis for the pan agonism of Rotigotine for the dopaminergic receptor system is unclear.

To date, several structures of dopamine receptors have been reported, including active state D1R, D2R, and D3R and inactive state D2R, D3R, and D4R ^11–16^. No active state structure of D4R or any state structure of D5R has been reported. The lack of the D5R structure and the active state D4R structure impedes our understanding on the dopaminergic system. In addition, the basis of which different types of dopamine receptors bind ligands with similar or diverse affinity is not well understood, making it is challenging to develop therapeutic agents with lower side effects. Here, we report cryo-electron microscopy (cryo-EM) structures of all five subtypes of dopamine receptors in complex with Rotigotine and their cognate G-protein subtypes of G_s_ and G_i_. These structures reveal the basis for the pan agonism of Rotigotine and dopamine receptor polypharmacology, as well as a mechanism of dopamine receptor activation and G-protein coupling selectivity. Together with mutagenesis and functional studies, our results provide important insights into the biology of dopaminergic system and templates for rational design of drugs treating CNS diseases.

## RESULTS AND DISCUSSION

### Cryo-EM structures of the five sub-type dopamine receptors

For cryo-EM studies, we used the wild-type human dopamine receptors for structural determination. To assist the expression and purification of the receptor-G protein complexes, we fused Bril ^17^ and His tags at the N-terminus of the receptors. To achieve the stable formation of the receptors with G-proteins, D1R or D5R were expressed with the dominant-negative form of G_s_ (DNG_s_) ^18^, and D2R or D3R were expressed with dominant-negative form G_i_ (DNG_i_) ^19^ in *Trichoplusia ni* insect cell. The D4R-G_i_ complex was not assembled stably, and the yield was low. To obtain the stable complex of D4R bound to G_i_-protein, we used another form of engineered G_i_-protein, which consists of αN and α5 helices of G_i_ and Ras domain of DNG_s_ ^20^. In addition, NanoBiT tethering strategy was introduced to enhance the assembly of the D4R-G_i_ complex by fusing the LgBiT to the C-terminus of the receptor and fusing the SmBiT to the C-terminus of Gβ ^21^. For the D1R-G_s_ and D5R-G_s_ complex, nanobody35 (Nb35) ^22^ were used to further stabilize the complexes. For the D2R-G_i_, D3R-G_i_, and D4R-G_i_ complexes, scFv16 ^23^ was used. The pan agonist Rotigotine and apyrase were added during purification to stabilize the complex in the active state. The structures were determined at a global resolution of 3.2 Å (D1R-G_s_), 3.0 Å (D2R-G_i_), 2.7 Å (D3R-G_i_), 3.1 Å (D4R-G_i_), and 3.1 Å (D5R-G_s_) (Fig. 1, Fig. S1, S2, Table S1). The density maps of the five complexes allowed us to model the majority of the receptor residues, ligands, and G-proteins, as well as a number of cholesterol molecules in D1R, D4R, and D5R (Fig. 1a, b, d). Several regions in the complexes were not observed in the EM maps, including the flexible N-terminus, a portion of extracellular loop 2 (ECL2), intracellular loop 3 (ICL3), C-termini of the receptors, and the alpha-helical domains (AHD) of Gα subunits (Fig. 1b). Although Nb35 was added during the purification of both D1R-Gs and D5R-Gs complexes, the density of Nb35 was not observed in the D5R-Gs complex (Fig. 1a, Fig. S2m, n).

**Fig. 1.**
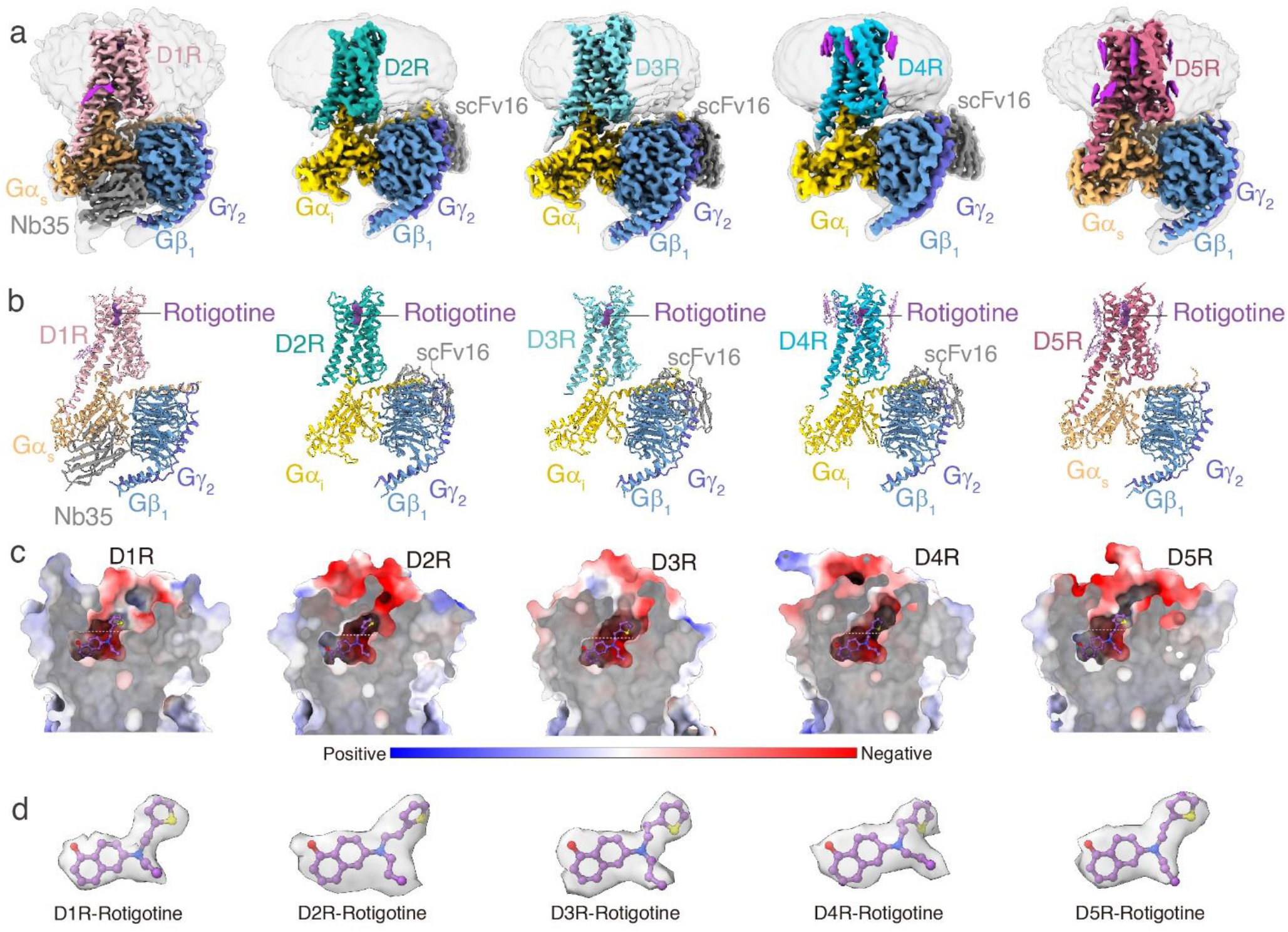
Cryo-EM structures of D1R, D2R, D3R, D4R and D5R signaling complexes. **a** The cryo-EM density maps of the D1R-Gs, D2R-Gi, D3R-Gi, D4R-Gi, and D5R-Gs complexes. **b** The models of the D1R-Gs, D2R-Gi, D3R-Gi, D4R-Gi, and D5R-Gs complexes. **c** The Ligand-binding pockets the D1R-Gs, D2R-Gi, D3R-Gi, D4R-Gi, and D5R-Gs complexes. **d** The Rotigotine structure in the D1R-Gs, D2R-Gi, D3R-Gi, D4R-Gi, and D5R-Gs complexes. The EM density of Rotigotine in the five structures is shown.

Overall, the five structures of the dopamine receptors exhibit similar backbone conformations (Fig. 2a). The seven transmembrane helical structures are highly overlapped, except for the extracellular side of TM1-3 (Fig. 2b-c), the intracellular side of TM5-6 (Fig. 2d), and ICL2 (Fig. 2e). Multiple cholesterol molecules were observed in the TMD of D1R, D4R, and D5R structures, but not in the D2R and D3R structures (Fig. 1a-e). The observation of cholesterols in D1R structure and the absence of cholesterols in D2R and D3R structures are consistence with the previously reported cryo-EM structures of D1R ^11,24^, D2R ^11^, and D3R ^12^. As the two members of D1-like receptor, D1R and D5R share almost identical backbone conformations, with RMSD of 0.48 Å by measured by the Cα atoms of the receptors (Fig. 2a). Our D1R-G_s_ complex is highly similar to the previous reported structures of D1R-Gs complexes solved by cryo-EM (Fig. S3a) ^11,24,25^. While the only x-ray structure of D1R-Gs complex^26^ shows a ~ 5 Å translation at αN helix of Gα_s_ subunit and Gβγ subunit from the cryo-EM structures (Fig. S3a). This suggests the crystal packing interactions on the αN helix may affect the conformation of the G-protein. For the D2-like group, D2R, D3R, and D4R also exhibited very similar conformation in their TMDs, but showed obvious differences in ECL2, ECL3, ICL2, and H8 helix (Fig. 2a). The D2R and D3R are more similar to each other than to D4R, with an RMSD of 0.51 Å between D2R and D3R, 0.73 Å between D2R and D4R, and 0.67 Å between D3R and D4R. This is consistent with the sequence identity between D2R and D3R (44%), which is higher than the sequence identity between D2R and D4R (31%) or between D3R and D4R (32%). The Rotigotine-bound D2R-G_i_ structure when compared with other cryo-EM structures of D2R-G_i_ protein complexes ^11,13^ showed conformational changes in both extracellular and intracellular regions, including TM6, TM7, ECL3, and G_i_ protein (Fig. S3c-e). In contrast, our D3R-G_i_ structure shares very high similarity with the previously reported D3R-G_i_ structures (Fig. S3f-g) ^12^. This may reflect that D2R is more dynamic in receptor conformation than D3R^12^. For the ligand-binding pocket, all five dopamine receptors exhibit highly negative charge properties, which allow the positively charged amine ligands to bind into their pockets (Fig. 1c). The differences between D1-like receptors and D2-like receptors were observed in the intracellular regions, including the intracellular end of TM5/6 and ICL2 (Fig. 2d-e). The most notable difference is in TM5 from the D1-like receptors, which is extended with three extra helical turns into the cytoplasmic side over the D2-like receptors (Fig. 2d). In addition, TM6 from the D1-like receptors are 7 Å more outward movement at the intracellular end than the D2-like receptors (Fig. 2d), consistent with their coupling selectivity of G_s_ and G_i_ subtypes^27^. For the ICL2 structure, the D1-like receptors exhibit two turns of α-helical while the D2-like receptor shows one turn of α-helical linked a loop (Fig. 2e).

**Fig. 2.**
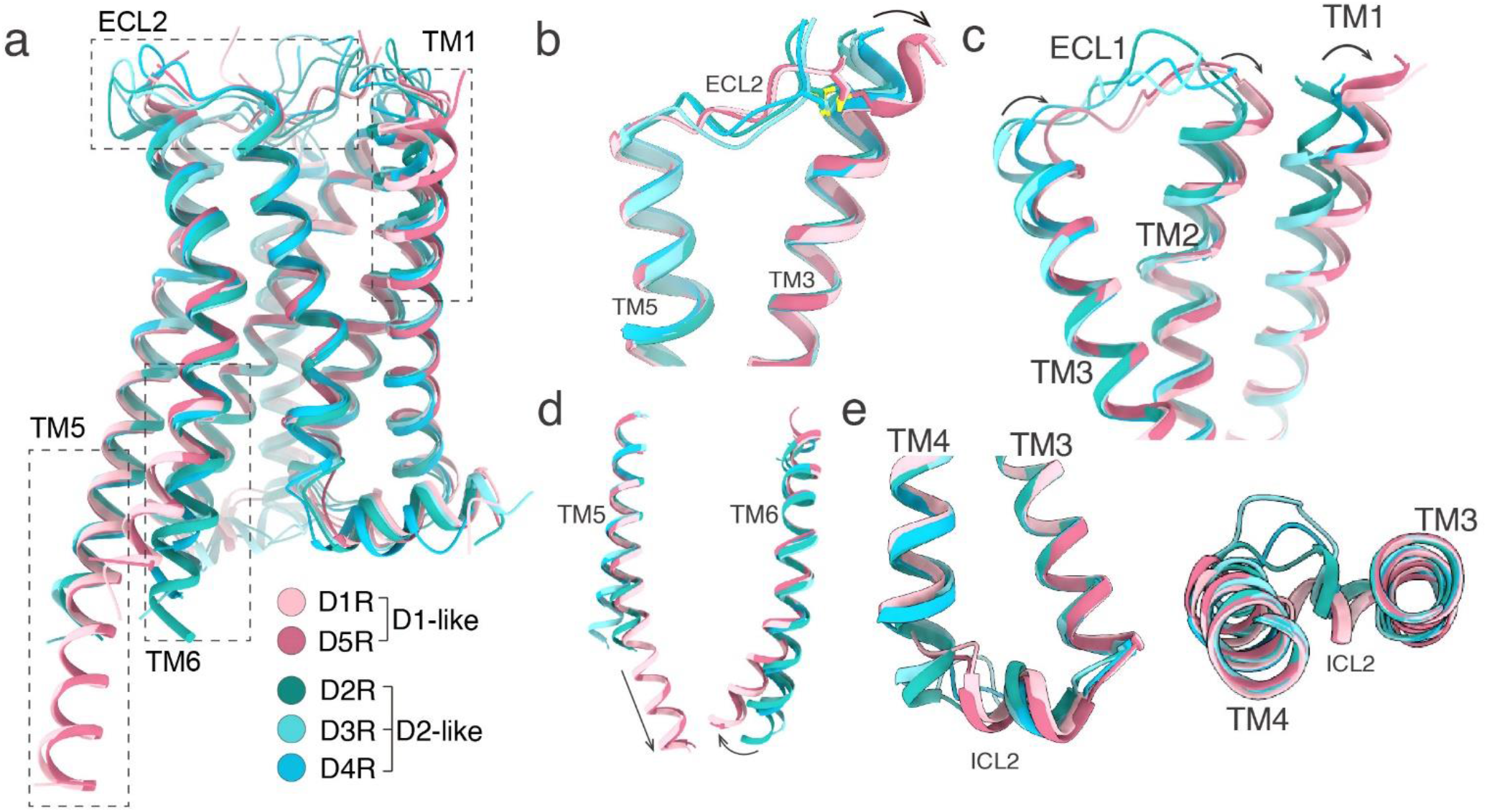
Structural features comparison of all active dopamine receptors. **a** Structural superposition of D1R, D2R, D3R, D4R, and D5R. **b** Structural alignment of ECL2, TM3, and TM5. **c** Structural alignment of ECL1 and TM1. **d** Structural alignment of TM5 and TM6. **e** Structural alignment of ICL2.

### Rotigotine binds to orthosteric and extended binding pockets

All five dopamine receptors harbor an open ligand binding pocket within the top half of their TMDs (Fig. 1c), which consists of an orthosteric binding pocket (OBP) and an extended binding pocket (EBP) (Fig. 1c). The OBP sits in the lower half of the entire pocket, reaching the middle of the receptor transmembrane helix. The EBP opens upward, connecting the OBP and the extracellular space (Fig. 1c). The sequences of OBPs share higher similarity than those of EBPs in five dopamine receptors, as well as in aminergic receptors^11,12^. Rotigotine binds to both OBPs and EBPs in all dopamine receptors (Fig. 1c and 3). In the OBPs, Rotigotine display a nearly identical conformation in all structures (Fig. 1c and 3). The primary amine group from Rotigotine forms charged interactions with the conserved D^3.32^ residue of the receptors, and the tetrahydronaphthalene group forms hydrophobic interactions with I^3.33^, F^6.51^ and F^6.52^ of the receptors (Fig. 3). The amino-linked ethyl group inserts into a small hydrophobic pocket formed by the receptor residues W^6.48^, W/Y^7.43^, and F^6.51^ (Fig. 3). In the EBPs, Rotigotine exhibits a conserved binding mode in D1R and D5R, while displays different binding modes among D2R, D3R, and D4R (Fig. 1d and 3). The hydroxyl group of Rotigotine forms a hydrogen bond to S^5.42^, which is a conserved residue in all dopamine receptors and forms another hydrogen bond to N^6.55^ of D1R and D5R (Fig. 3). However, the S^5.42^ hydroxyl group shows different interaction patterns in D2-like receptors with the hydroxyl group of Rotigotine only forms hydrogen bond to H^6.55^ in D3R and D4R, but not in D2R, although the H^6.55^ residue is conserved in D2-like receptors (Fig. 3b-d). Furthermore, the D3R residue H^6.55^ is stabilized by residue Y^7.35^ through a hydrogen bond (Fig. 3c). The 7.35 position in D4R is V^7.35^ and it cannot form the corresponding hydrogen bond with H^6.55^ in D4R suggesting that the binding of Rotigotine to D3R is stronger than D4R due to the interactions with H^6.55^ in D3R but not in D4R. For the comparison of D2R and D3R, despite the H^6.55^ and Y^7.35^ residues are conserved and the H^6.55^-Y^7.35^ hydrogen bond could be formed in both receptors, the extracellular end of TM6 and TM7 in D2R is moved outward relative to D3R (Fig. 3). This conformational difference makes the H^6.55^ residue in D2R is far away to form a hydrogen bond with the hydroxyl group of Rotigotine (Fig. 3). Thus, the different patterns of the interactions between the hydroxyl group of Rotigotine and D2-like receptors suggests the basis for D3R to have higher affinity than D2R and D4R, consistent with our functional experiments (Table S2–5). The thiophene group of Rotigotine forms hydrophobic interactions with residues from the EBPs of the five dopamine receptors (Fig. 1c and 3). In both D1R and D5R structures, the thiophene group shares a nearly identical interaction mode between the two receptors because of conserved residues and conformations in the EBPs (Fig. 3a, e, Fig. S4), where the thiophene group forms hydrophobic interactions with receptor residues W^3.28^, F^7.35^, and V^7.39^ (Fig. 3a, e). On the other hand, in the structures of the D2-like receptors, the thiophene group of Rotigotine displays different conformations due to the different shapes and topology of their EBPs (Fig. 3b-d, Fig. S4). The D2R and D3R structures show that the thiophene group forms hydrophobic interactions with residues F^3.28^, V^2.61^, T^7.39^, Y^7.35^, and I/S^ECL2^. Interestingly, the thiophene group displays a different interaction mode in D4R from D2R and D3R. In D4R, the thiophene group is pointed toward to TM2 and forms hydrophobic interactions with residue F91^2.61^ (Fig. 3d), which is not conserved in other dopamine receptors. The corresponding residue is L^2.61^ in D1R/D5R and V^2.61^ in D2R/D3R. The affinity of Rotigotine for D4R was reduced by 100 folds with the mutation F91A^2.61^, whereas the corresponding alanine mutation of residues 2.61 at other dopamine receptors had little effect on Rotigotine binding (Fig. 3, Table S4). These results support the unique interacting mode of Rotigotine in D4R.

**Fig. 3.**
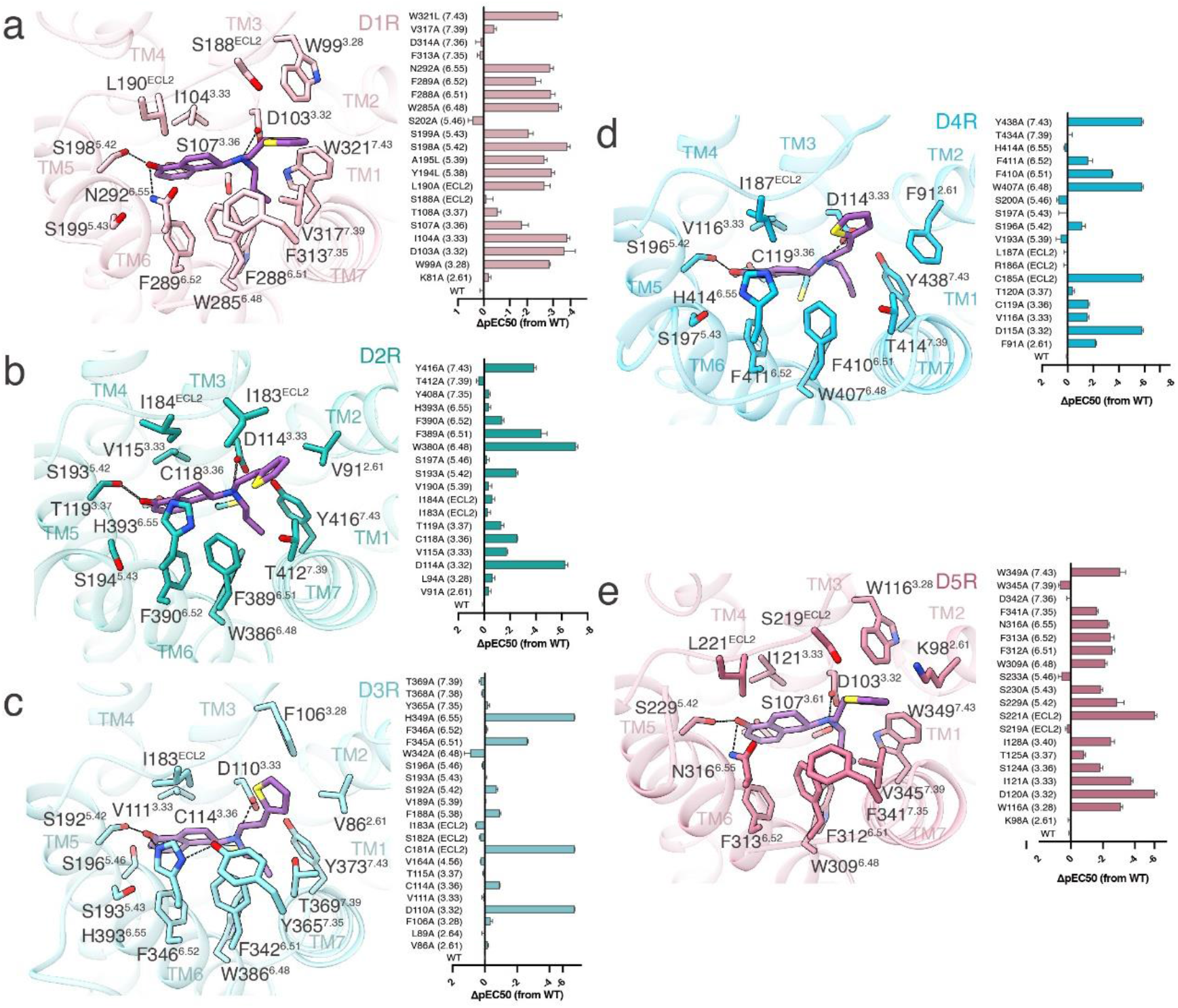
Rotigotine recognition at all dopamine receptors. **a** Detailed interaction between Rotigotine and D1R (left). Effects of mutations of the ligand-binding pocket residues of D1R on changes in ΔpEC_50_ in response to stimulation of Rotigotine, evaluated using a GloSensor cAMP assay (right). **b** Detailed interaction between Rotigotine and D2R (left). Effects of mutations of the ligand-binding pocket residues of D2R on changes in ΔpEC_50_ in response to stimulation of Rotigotine, evaluated using a GloSensor cAMP assay (right). **c** Detailed interaction between Rotigotine and D3R (left). Effects of mutations of the ligand-binding pocket residues of D3R on changes in ΔpEC_50_ in response to stimulation of Rotigotine, evaluated using a GloSensor cAMP assay (right). **d** Detailed interaction between Rotigotine and D4R (left). Effects of mutations of the ligand-binding pocket residues of D4R on changes in ΔpEC_50_ in response to stimulation of Rotigotine, evaluated using a GloSensor cAMP assay (right). **e** Detailed interaction between Rotigotine and D5R (left). Effects of mutations of the ligand-binding pocket residues of D5R on changes in ΔpEC_50_ in response to stimulation of Rotigotine, evaluated using a GloSensor cAMP assay (right). All data are presented as mean values ± SEM of three independent experiments for the wild type (WT) and mutants (n=3).

Since the OBP sequences of dopamine receptors are highly conserved, many ligands were developed using the same chemical scaffold. Catechol and ergoline compounds are two classes of prototypical ligands of dopamine receptors and other aminergic receptors. Rotigotine does not belongs to either catechol or ergoline class of ligands. To uncover the differences of the Rotigotine binding modes from the catechol agonists, we analyzed the detailed interactions by comparing the structures of dopamine receptors bound to Rotigotine and dopamine (a prototypical catechol ligand)^25^, which reveals five sets of intermolecular interactions between the bound ligands and the receptors (Fig. S4c-e). Among the five set of interactions, three sets are similar, and two sets are different (Fig. S4c-e). The three similar sets of interactions include the conserved salt bridge formed by the primary amine group with the D^3.32^ residue, the hydrophobic interactions of the tetrahydronaphthalene group of Rotigotine and the benzene group of dopamine with hydrophobic residues 6.51, 6.52, and 3.33, and the hydrogen bond interactions of the 5-hydroxyl group of Rotigotine and dopamine with polar residues S^5.42^, S^5.43^, and N^6.55^ (Fig. S4c-d). Mutations of these residues to Ala significantly reduced the potencies of both Rotigotine and dopamine to D1R by similar degrees (Tables S2). The two different interactions include extra hydrogen bonds of the 4-hydroxyl group from dopamine with residues S202 and T108 while the corresponding interactions cannot be formed by Rotigotine because Rotigotine does not have the corresponding hydroxyl group as in dopamine (Fig. S4c-e). Correspondingly, S202A or T108A mutations in all dopamine receptors reduced the dopamine potencies by over 1000 folds, but hardly affected the potencies of Rotigotine (S202A) or mildly reduced the potencies of Rotigotine by 20 folds for all dopamine receptors (T108A) (Tables S2). The second set of different interactions is seen in the EBPs because dopamine lacks a branch group like the thiophene group in Rotigotine, which forms additional hydrophobic contacts with the EBPs (Fig. S4c-e). Correspondingly, mutations on EBP residues in all five dopamine receptors affected the potencies of Rotigotine more significantly than those of dopamine, which is mostly bound within the OBP (Fig. Tables S2).

### Rotigotine Polypharmacology

Rotigotine has been reported to have the ability to activate all dopamine receptors, as well as other several types of aminergic receptors ^28^. To reveal the polypharmacological profile of Rotigotine, we screened the binding activity of Rotigotine to over 300 GPCRs (Table S7). The results showed that Rotigotine exhibited high affinities to dopamine receptors, serotonin receptors, and adrenergic receptors (Fig. 4, Table S7) and unexpectedly displayed agonist activities at somatostatin receptors, adenosine receptors, opioid receptors, and melatonin receptors (Table S7). To illustrate the basis of the promiscuous binding for Rotigotine, we aligned the sequence of the ligand-binding pocket of aminergic receptors (Fig. 4d). We found that the high affinity of Rotigotine is highly related to the conserved sequences of OBP in many monoamine receptors (Fig. 4d). Structure comparisons of the Rotigotine-bound dopamine receptors with serotonin receptors and adrenergic receptors revealed highly overlapped conformations shared by the conserved OBP residues, including D^3.32^, I/V^3.33^, F^6.51^, F^6.52^, and W^6.48^ (Fig. 4e-g). Mutations in these conserved OBP residues in dopamine receptors greatly affect Rotigotine binding (Table S2–6), indicating that the polypharmacology of Rotigotine to many GPCRs is mainly contributed by the conserved OBP.

**Fig. 4.**
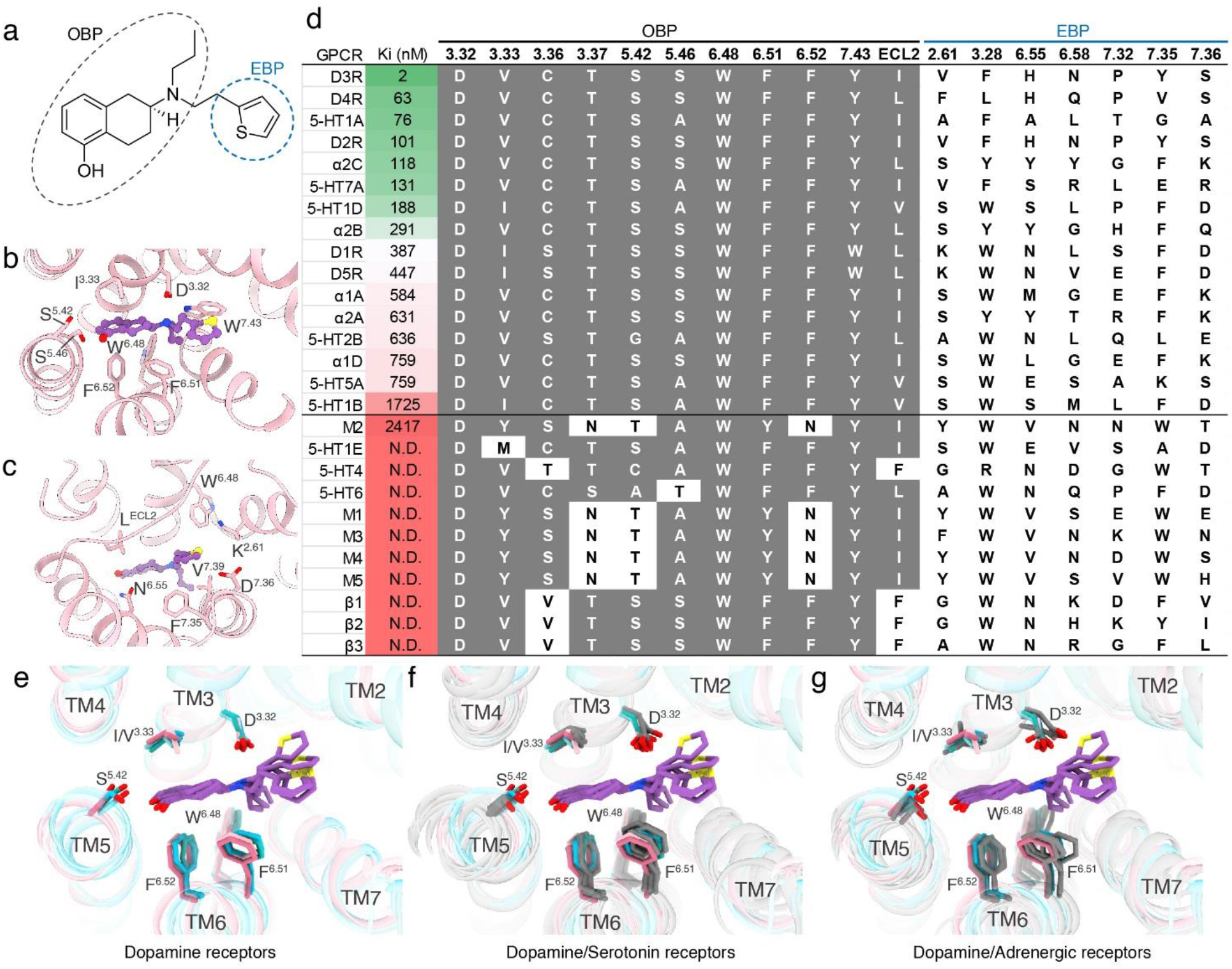
Polypharmacological profile of Rotigotine. **a** The chemical structure of Rotigotine. **b** The interaction of D1R OBP with Rotigotine. **c** The interaction of D1R EBP with Rotigotine. **d** Rotigotine affinities (Ki) from radioligand competition binding assays and the OBP-EBP residues alignment. Receptors are listed in order of decreasing Rotigotine affinity. **e** Structural superposition of five dopamine receptors and bound Rotigotine. **f** Structural superposition of five dopamine receptors, bound Rotigotine in compared with serotonin receptors (gray): 5-HT_1A_ (7E2Y), 5-HT_1B_ (6G79), 5-HT_1D_ (7E32), 5-HT_2B_ (6DRY), and 5-HT5A (7X5H). **g** Structural superposition of five dopamine receptors, bound Rotigotine in compared with adrenergic receptors (gray): a2A (6KUY), a2B (6K41), and a2C (6KUW).

### D1-like receptors

The two D1-like dopamine receptors, D1R and D5R, exhibit highly conserved sequence homology, particularly at the orthosteric binding pocket (Fig. 4d). To further explore the remaining differences in ligand affinity for D1R and D5R, we performed structural superimposition of Rotigotine-bound receptors. The Rotigotine-bound D1R and D5R structures revealed that they share almost identical conformations in their ligand-binding pockets (Fig. 5a-c). However, both dopamine and Rotigotine have a higher potency to D5R over D1R (Fig. S4a, Table S2, S6). The cAMP accumulation assays showed that the potency of dopamine is approximately 10-fold more potent on D5R (pEC_50_ = 9.82) than on D1R (pEC_50_ = 8.86) and Rotigotine is also ~10-fold more potent on D5R (pEC_50_ = 9.25) than on D1R (pEC_50_ = 8.49) (Table S2, S6). A comparison of their ligand-binding pockets and their electrostatic surface show that D5R has more negative charges on the extracellular surface than D1R (Fig. 5d-e). These findings indicate that positively charged ligands, such as dopamine and Rotigotine, would prefer to be enriched within the pocket of D5R. In addition, we performed a docking study of Rotigotine into the ligand-binding pocket of D1R and D5R, which revealed that Rotigotine had a better docking score on D5R (−7.09) than on D1R (−4.85), consistent with our functional studies. Further comparison of the residue pair F^7.35^-L^6.58^ of D1R with the F^7.35^-V^6.58^ of D5R one can find a subtle difference in the Rotigotine binding modes between these two receptors. Residue F^7.35^ in EBPs forms hydrophobic interactions with the thiophene group of Rotigotine in both D1R and D5R (Fig. 5f). However, the side chain of F^7.35^ in D5R is slightly closer to the thiophene group of Rotigotine than in D1R (Fig. 5f). Thus, the F341^7.35^A mutation in D5R had a greater effect on Rotigotine binding (ΔpEC_50_=-1.63) than the corresponding F313^7.35^A mutation in D1R (ΔpEC_50_=0.16) (Table S2, S6).

**Fig. 5.**
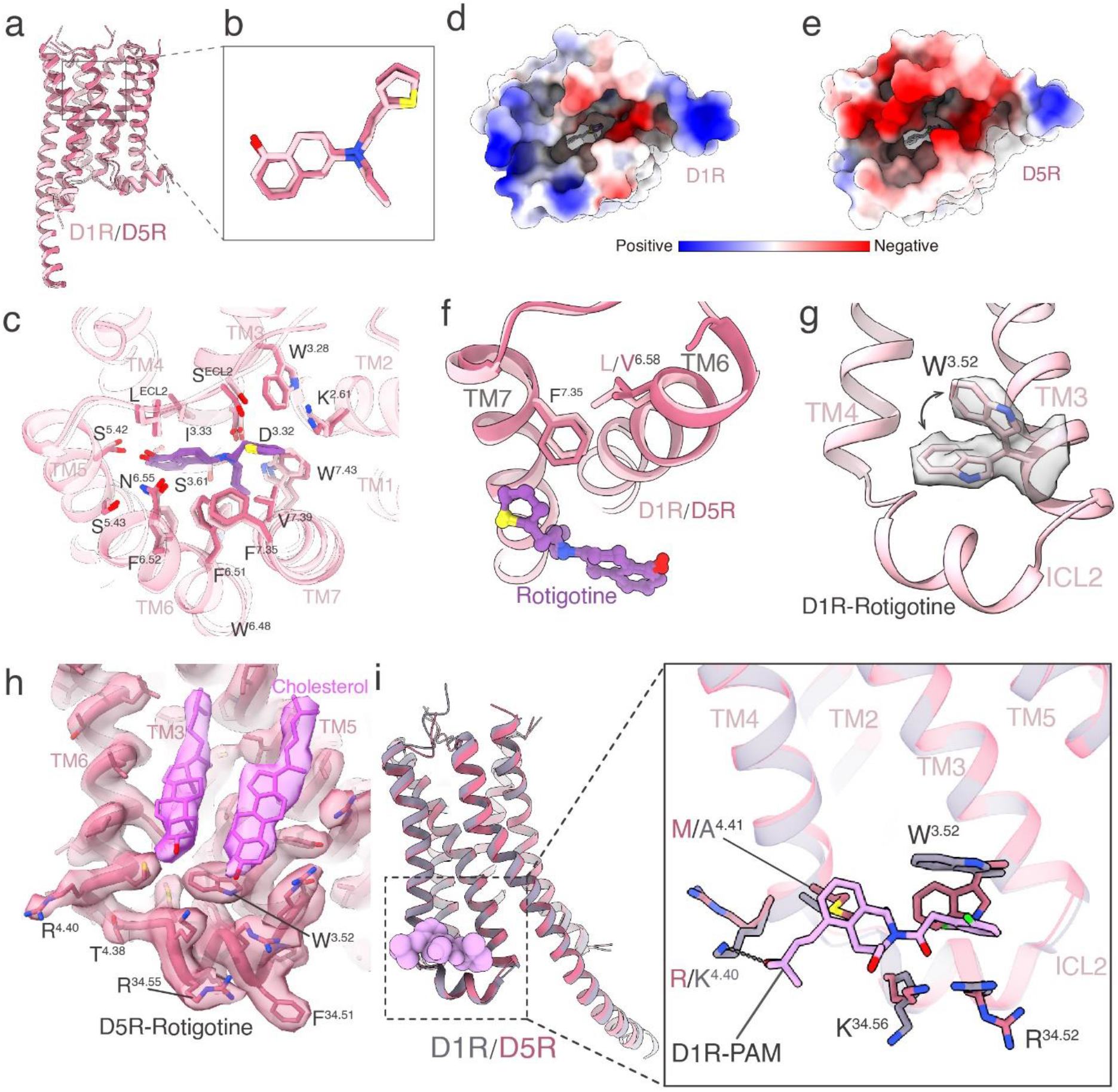
Comparison of D1R and D5R in Rotigotine-binding and PAM-binding. **a** Structural superposition of D1R-G_s_ and D5R-G_s_ complex when receptors were aligned. **b** Comparison of Rotigotine binding poses of D1R and D5R structures. **c** Structural comparison of Rotigotine recognition between D1R and D5R. **d-e** Rotigotine-binding pockets of D1R (**d**) and D5R (**e**) viewed from the extracellular side. **f** Comparison of TM6 and TM7 residues for Rotigotine recognition between D1R and D5R. **g** The sidechain of W^3.52^ residue shows alternative conformation in the D1R-Gs-Rotigotine structure. **h** The unique conformation of W^3.52^ residue is stabilized by a cholesterol molecule in the D5R-Gs-Rotigotine structure. **i** Comparison of TM3, TM4 and ICL2 residues for compound LY3154207 (a high selective PAM for D1R) recognition between D1R and D5R.

In addition to orthosteric agonists, positive allosteric modulators (PAM) represent a promising strategy for drug discovery of D1R and D5R with high selectivity and low side effect. LY3154207 is the first clinical PAM for D1R with a high level of selectivity^29^. To explore the PAM selectivity between D1R and D5R, we compared the structures of the LY3154207-bound D1R with the Rotigotine-bound D1R and D5R. A notable difference was that A^4.41^ in D1R was replaced by M^4.41^ in D5R, which result in a steric hindrance for D5R to bind LY3154207 (Fig. 5i). In addition, W^3.52^ of D1R was found to have two alternative conformations in the Rotigotine-bound D1R structure (Fig. 5g). One of the conformations, with the side chain in the up configuration, could adapt to the LY3154207 binding^25^. The other conformation, with the side chain in the down configuration, would prevent LY3154207 from binding (Fig. 5g, i). In the D5R structure, residue W^3.52^ only has one conformation in its down configuration, which would prevent D5R from binding to LY3154207 (Fig. 5h). The unique conformation of W^3.52^ in D5R is further stabilized by a cholesterol molecule (Fig. 5h), which is not observed in the corresponding site of D1R. Thus, the structures determined here provide a framework for understanding the mechanism of PAM selectivity and could assist in the design and optimization of D1R selective therapeutic modulators.

### D2-like receptors

Despite that the three D2-like receptors share relatively high sequence conservation and all of them couple to G_i_ protein, they play different physiological functions and show different affinities for various ligands^4^. To reveal the structural differences between D2R and D3R in the active state, we compared the Rotigotine-bound D2R and D3R structures with the previously reported active D2R and D3R structure^11,12^. The comparison shows obvious conformational changes in the extracellular end of TM6/7 between the structures of Rotigotine- and Bromocriptine-bound D2R, especially at ECL2 (Fig. S3c-e). However, in contrast to variations between the D2R structures, comparison of D3R structures show almost identical conformation in all TMD structural elements (Fig. S3f-g). This is consistent with the notion that D2R is more dynamic than D3R ^12^. Despite no other active state D4R structure available now, we compared the structures of D4R with D2R and D3R. We found an apparent difference between D4R and D2/3R in that there were multiple cholesterol molecules surrounding the D4R TMD but not in D2R and D3R (Fig. 1a). Remarkably, a clear cholesterol molecule is located between TM1 and TM7 in D4R (Fig. 6a-b). This cholesterol forms hydrophobic interactions with W387^7.40^, which is further stabilized by the pocket residue F91^2.61^ (Fig. 6b). Interestingly, this cholesterol is only found in the D4R structure and the 2.61 residue is not conserved in all other dopamine receptors (Fig. 6b-f), mutation of residue 2.61 to Ala significantly reduced the potency of Rotigotine for D4R but not for other dopamine receptors (Tables S2–6), suggesting that the CHL-W^7.40^-F91^2.61^ interaction network is important for ligand binding in D4R. A similar cholesterol interacting network has also been observed in the 5-HT_1A_ receptor-ligand complexes^30^.

**Fig. 6.**
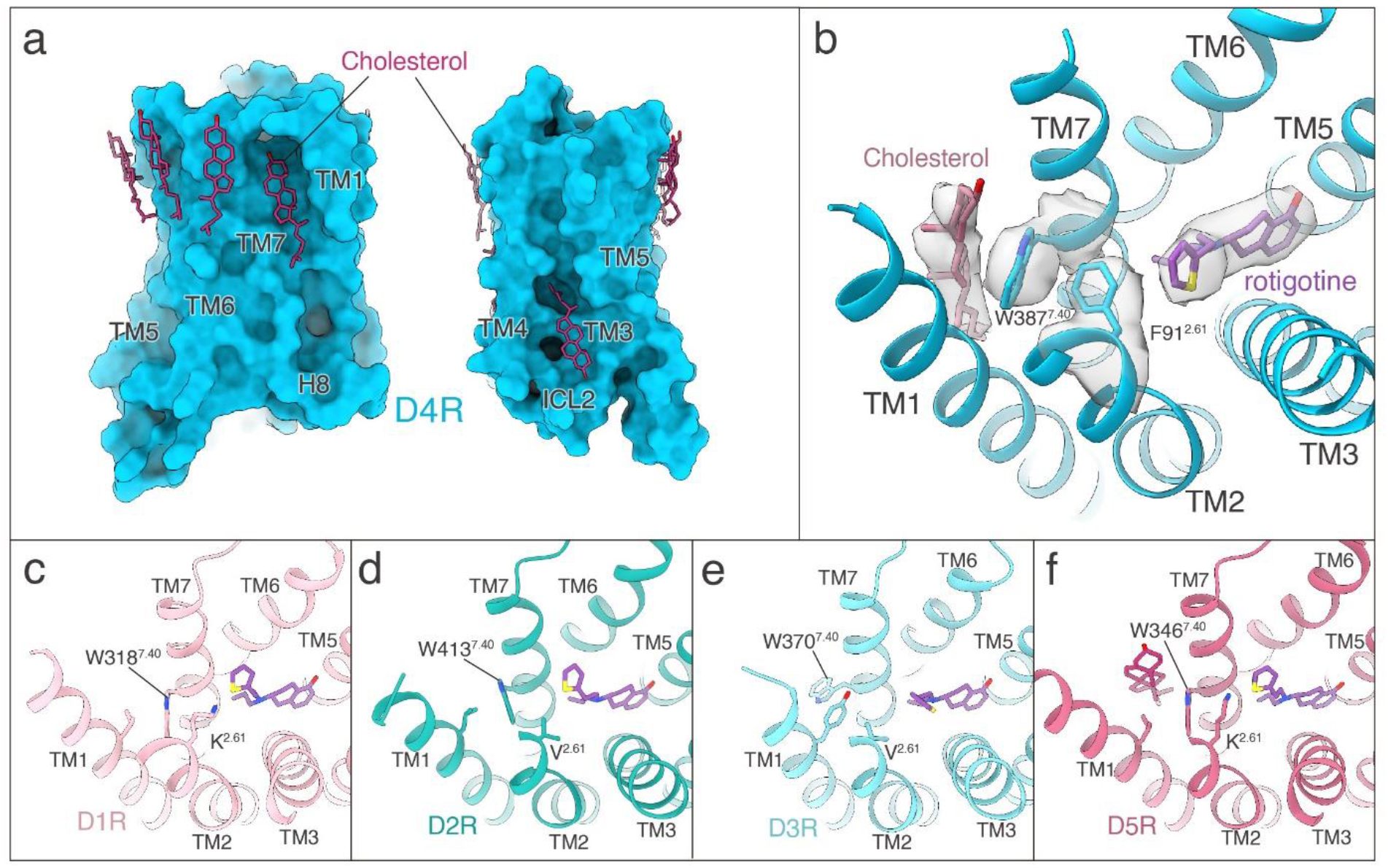
The binding of Rotigotine in D4R is regulated by cholesterol. **a** Cholesterol molecules at the surface of D4R. **b** A cholesterol molecule is located between TM1 and TM7 of D4R and stabilizes Rotigotine binding through residues W387^7.40^ and F91^2.61^. **c-f** The comparison of the TM1-TM7 structure and residue 2.61 from D1R (**c**), D2R (**d**), D3R (**e**), and D5R (**f**) show difference from D4R.

### Activation of the Dopamine receptors

The availability of all five dopamine receptor structures allowed us to examine the common features of dopamine receptor activation mechanism. The activation of D4 and D5 receptors has similar characteristics to the previously reported activation of D1R, D2R, and D3R^11,12^. For all five dopamine receptors, the binding of Rotigotine pushes the “toggle switch” residue W^6.48^ downward, which further induces conformational changes in the PIF, DRY and NPxxY motifs, which eventually cause the outward movement of TM6, allowing the α5 helical of G-protein to insert into the intracellular pocket of the TMD. Within the D1-like receptors, the active structures of both D1R and D5R shares nearly identical conformations in the “toggle switch” residue, PIF, DRY, and NPxxY motifs, suggesting a common mechanism of Rotigotine-induced activation shared by both D1R and D5R. Within the D2-like receptors, all motifs related to receptor activation are conserved between D2R and D3R, and the motif residues shares very similar conformational changes between the inactive and active structures (Fig. S5). However, the D4R structure shows different conformational changes from D2R and D3R (Fig. S5). In particular, the intracellular end of TM3 in the active D4R undergoes a 3 Å inward translation from the inactive state (Fig. S5). This translation is not observed between the active and inactive structures of D2R and D3R (Fig. S5), revealing a unique feature of D4R activation.

Previous studies reported that the catechol agonists can activate the aminergic receptor through a hydrogen bond with S^5.46^, which induces an inward movement of residue P^5.50^ and the rearrangement of the PIF motif and TM6 ^31,32^. In all dopamine receptor structures, Rotigotine does not form the same hydrogen bond with the S^5.46^ as dopamine (a catechol ligand with two hydroxyl groups) because Rotigotine contains only one hydroxyl group, which form hydrogen bonds with S^5.42^ and N^6.55^ (Fig. S4c-d). However, Rotigotine, despite not forming hydrogen bond with S^5.46^, can still activate dopamine receptors with similar efficiency as dopamine (Table S2–6). To reveal the different activation mechanisms of Rotigotine and catechol agonists, we compared the Rotigotine-bound D1R structure with the dopamine-bound D1R structure^11^. We found that the closest distance between residue W^6.48^ and Rotigotine is 3.4 Å, but for dopamine this distance is 5.3 Å (Table S8). This distance difference may cause the different strength of interactions of Rotigotine and dopamine with W^6.48^, therefore affecting their activation of D1R. Consistent with these observations, mutation W^6.48^A in D1R has a greater reduction of G protein signaling for Rotigotine than dopamine, although Rotigotine exhibit a higher affinity for the mutated receptor (Table S2 and S8). These results suggest that dopamine activates receptors through both the residues S^5.46^ and W^6.48^, whereas Rotigotine activates receptors mainly through residue W^6.48^. Our results indicate that there are different activation mechanisms of catechol and non-catechol agonists for dopamine receptors.

To illustrate the agonism and antagonism of dopamine receptors, we analyzed all available dopamine receptor structures and focused on the interactions of Rotigotine with residue W^6.48^, as well as the interactions of residue W^6.48^ with the PIF motif. We found the activation of the respective receptor is highly related to the distance between residue W^6.48^ and I^3.40^. Specifically, all activated dopamine receptors have closer distances (< 5 Å) of the W^6.48^-I^3.40^ distance than all inactive dopamine receptors (> 6 Å). Except for the antagonist L745870-bound D4R structure, the W^6.48^-I^3.40^ distance is 4.9 Å, but the structural activation analysis shows that this antagonist-bound D4R structure exhibits 83% activation ^33^. The typical antagonist of dopamine receptors could prevent the W^6.48^-I^3.40^ contacts by inserting deeply into the OBP, whereas agonists only bind to the upper half of receptors. These results suggest the importance of W^6.48^-I^3.40^ interaction for receptor activation and further reveal the basis of agonism and antagonism of dopamine receptors.

### G-protein Coupling of Dopamine Receptors

The interactions of the five subtypes of dopamine receptors with the respective G-protein displays a relatively conserved mode as other aminergic receptors, with remarkable features that fit the TM5-TM6 switches for G_s_ and G_i/o_ selectivity^27^. For the G_s_-coupled D1R and D5R, their TM5s are extended into the cytoplasmic side that end at residue 5.84, and form extensive interactions with the Gαs-Ras domain (Fig. 7a). In contrast, for the G_i_-coupled D2R, D3R and D4R, their TM5s are not extended as those of D1R and D5R, which end at residue as 5.69 (D2R and D4R) or 5.73 (D3R) (Fig. 7a). The TM5-Ras interactions were not formed by the D2-like receptors with their G_i_ protein (Fig. 7a). This Gi- and Gs-coupling selectivity is consistent with the determinants of TM5-TM6 switches for the G_i_ and G_s_ selectivity as originally revealed by serotonin receptors^27^. In addition to the differences in the receptor structures, several differences were also observed in the orientation of the G proteins between G_i_ and G_s_ complexes (Fig. 7), with the α5 helices of Gαs subunit showing a 4.5 Å translation from the G_i_ complexes (Fig. 7c). At the ICL2 of receptors, the 34.51 residue is conserved as a hydrophobic residue and forms hydrophobic interactions with G-protein by inserting its sidechain into the cleft between αN and α5 of G-protein (Fig. S7a-f). The hydrophobic interactions between 34.51 and Gα cleft are conserved in all dopamine receptors and many other GPCRs^30,34–40^. On the other hand, the rest of ICL2, which is not conserved in sequence, forms different interactions with the G proteins among the five dopamine receptors (Fig. S7). Specifically, D1R residue E132^34.54^ forms unique polar interactions with G_s_ residue H41 (Fig. S7b), D2R residue Y146^34.57^ forms unique polar interactions with G_i_ residue E28 (Fig. S7d), and D3R residue H140^34.55^ forms unique polar interactions with the main chain of G_i_ residue A31 (Fig. S7e). Together, these structural observations reveal common and unique features that determine G protein coupling specificity of dopamine receptors.

**Fig. 7.**
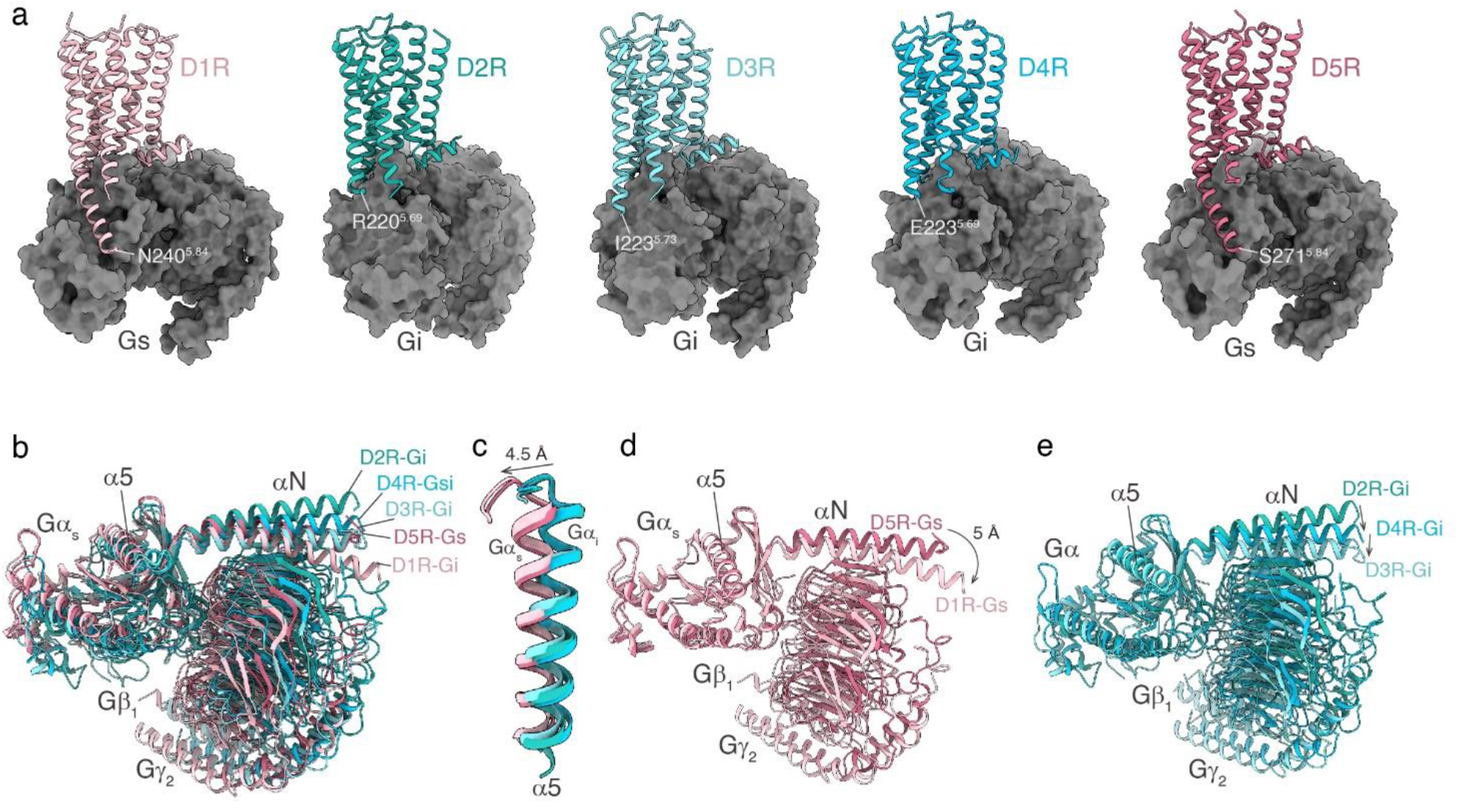
G-protein coupling of dopamine receptors. **a** The structures of the D1R-G_s_, D2R-G_i_, D3R-G_i_, D4R-G_i_, and D5R-G_s_ complex. **b** Comparison of the G proteins conformation among the structures of D1R-G_s_, D2R-G_i_, D3R-G_i_, D4R-G_i_, and D5R-G_s_ complex. **c** Structure comparison focuses on the α5 helix of the Gα subunit bound to dopamine receptors. **d** Comparison of the G proteins conformation among the structures of D1R-G_s_ and D5R-G_s_ complex. **e** Comparison of the G proteins conformation among the structures of D2R-G_i_, D3R-G_i_, and D4R-G_i_ complex.

### Concluding Remarks

Here, we report the cryo-EM structures of all five dopamine receptors, D1R, D2R, D3R, D4R, and D5R, in complex with G-proteins, among which the D4R and D5R structures are the first set of their active structures. The structures reveal a universal binding mode of the pan agonist Rotigotine in all five dopamine receptors and the specific intermolecular interactions that define the recognition of Rotigotine by each of dopamine receptors. Structural and sequence comparisons indicate that the conserved OBP is the basis for promiscuous binding of Rotigotine with all five dopamine receptors as well as with many other monoamine receptors, including receptors subtypes for serotonin, adrenergic amines, histamine, and muscarinic amines. Rotigotine is mainly prescribed for PD and restless legs syndrome (RLS), with potential anti-depression effects from its cross reactivity of serotonin receptors 5-HT_1A_ (Fig. 4d). The structures and the binding results from this study therefore provide a rational basis for understanding the profound polypharmacology of Rotigotine and its therapeutic effects.

The structures of all five dopamine receptors reveal a highly similar OBP where Rotigotine binds, thus posing a great challenge to the design of subtype specific orthosteric agonists. As an adjunct, one alternative strategy is to design subtype specific allosteric modulators, such as LY315420, which is a D1R specific PAM. Our structures reveal that D1R residues A^4.41^ and W^3.52^ are key to the selective PAM agonism of LY3154207 on D1R. These structural observations provide critical insights into the PAM selectivity of LY315420 for D1R over D5R, and the basis for designing next generations PAMs targeting dopamine receptors.

The five dopamine receptor structures also reveal differential roles of cholesterol in dopamine signaling. Specifically, a number of cholesterol molecules are found in D1R, D4R and D5R, but not in D2R and D3R. Most of these cholesterol molecules are found to surround the extracellular half of the TMDs, in analogous to cholesterol molecules in the structures of 5-HT_1A_^30^ and class B GPCRs such as CRFR1, CRFR2 ^41^, and PTH1R^42^. The regulatory roles of cholesterol have been showed to be important in 5-HT_1A_^30^ and CRFR1/2 ^41^. In this paper, we also showed that cholesterol is involved in ligand binding. For example, the D4R structure reveals the mechanism of cholesterol in the regulation of ligand binding in D4R through an interaction network formed by a cholesterol at the cleft of TM1 and TM7 (Fig. 6b). The location of this cholesterol in D4R is nearly identical to that in 5-HT_1A_, suggesting the conserved role of cholesterol in ligand binding of different monoamine GPCRs.

Through structural comparisons, we also uncovered conserved activation mechanisms of dopamine receptors and the detailed conformational changes during activation, as well as the basis of agonism and antagonism. We have additionally analyzed the selectivity and the unique features of all five dopamine receptors in G-protein coupling. Together, our work presents the structural genomics of the human dopamine receptor system and provides structural templates for the development of selective or non-selective agonists, antagonists, and allosteric modulators of dopamine receptors, with potential significance for the treatment of CNS diseases.

## ACKNOWLEDGMENTS

The cryo-EM data were collected at the Center of Cryo-Electron Microscopy, Shanghai Institute of Materia Medica, and the Center of Cryo-Electron Microscopy, Zhejiang University. This work was partially supported by the National Key R&D Programs of China 2018YFA0507002, Shanghai Municipal Science and Technology Major Project 2019SHZDZX02 and XDB08020303 to H.E.X.; the National Key Basic Research Program of China 2019YFA0508800, National Natural Science Foundation of China 81922071, Zhejiang Province Natural Science Fund for Excellent Young Scholars LR19H310001 and Fundamental Research Funds for the Central Universities 2019XZZX001-01-06 to Y.Z.; National Natural Science Foundation 31770796 and National Science and Technology Major Project 2018ZX09711002 to Y.J.; grants from the NIMH Psychoactive Drug Screening Program (X-P. H., Y.L., B.L.R.) and RO1MH112205 to B.E.K. and B.L.R.

## AUTHOR CONTRIBUTIONS

P.X. and S.H. designed the expression constructs, purified the complexes, and prepared protein samples for the D2R-G_i_, D3R-G_i_, D4R-G_i_, and D5R-G_s_ complexes for cryo-EM data collection, performed cryo-EM grids preparation, data acquisition, and structure determination, and prepared the draft of the manuscript and Figures. Y.Z. (Youwen Zhuang) designed the constructs, prepared the protein sample, conducted cryo-EM data collection and structure determination of D1R-G_s_, and participated in the preparation of supplementary Figures. C.M. screened the cryo-EM conditions, prepared the cryo-EM grids, collected cryo-EM images, processed the EM data of the D2R-G_i_ complex, and participated in the preparation of supplementary Figures. P.X. built the models and refined the structures. Y.Z. (Yumu Zhang) and H.L. participated in the sample preparation and screening of the D4R-G_i_ and the D5R-G_s_ complexes. Y.W. participated in the sample preparation and screening of the D1R-G_s_ complex. B.E.K., X-P. H., and Y.L. performed cAMP, GPCRome, Tango Assays, and radioligand binding assays. B.E.K. compiled assay data and participated in the preparation of the manuscript. X.H. performed the docking studies. W.Y. designed the Gα constructs used for D4R-G_i_ complex. Y.J. participated in the funding acquisition. Y.Z. (Yan Zhang) supervised C.M. and participated in manuscript editing. B.L.R. supervised pharmacological and mutagenesis experiments and participated in manuscript writing. H.E.X. conceived and supervised the project and wrote the manuscript with P.X.

## DECLARATION OF INTERESTS

The authors declare no competing interests.

## MATERIALS AND METHODS

### Data and Code Availability

The atomic coordinates and cryo-EM maps included in this study will be deposited in the Protein Data Bank and Electron Microscopy Data Bank, respectively. PDB: and EMDB: (D1R-G_s_-Nb35-Rotigotine), PDB: and EMDB: (D2R-G_i_-scFv16-Rotigotine), PDB: and EMDB: (D3R-G_i_-scFv16-Rotigotine), PDB: and EMDB: (D4R-G_s_-scFv16-Rotigotine), and PDB: and EMDB: (D5R-G_s_-Nb35-Rotigotine).

### Cell Lines

*Trichoplusia ni* (Hi5) cells were grown in ESF 921 medium at 27°C and 120 rpm. HEKT cells were grown in a humidified 37°C incubator with 5% CO_2_ using media supplemented with 100 I.U./mL penicillin and 100mg/mL streptomycin (Invitrogen). The human cell lines HEK293T were maintained in DMEM (VWR) containing 10% fetal bovine serum (FBS, VWR).

### Constructs

The human wild-type D1R, D2R, D3R, D4R, or D5R was cloned into the pFastBac (Thermo Fisher) vector using ClonExpress II One Step Cloning Kit (Vazyme Biotech Co., Ltd). An N-terminal haemagglutinin (HA) signal sequence followed by a Flag tag and a His-tag was fused with the receptor proteins to facilitate expression and purification. A fragment of b_2_AR N-terminal tail region was fused in D1R and Bril was fused in D2R, D3R, D4R, and D5R as the fusion proteins. For the G proteins, dominant-negative (DN) mutations were induced in Gα subunits to decrease the affinity of nucleotide binding to the heterotrimer Gαβγ complex. For the D1R-G_s_ and D5R-G_s_ complexes, a mini-G format of Gα_s_ was used. The mini-Gα_s_ was generated by deleting the alpha-helical domain of Gα_s_ and introducing stabilized mutations under the previously reported sequence ^43,44^. Two dominant-negative mutations G226A and A366S were also included into the mini-Gα_s_ ^45^. For the D2R-G_i_ and D3R-G_i_ complexes, a dominant-negative form of Gα_i_ (DNGα_i_) was constructed by site-directed mutagenesis to incorporate mutations S47N, G203A, E245A, and A326S ^46^. For the D4R-G_i_ complex, a form of Gα_si_ construct with α5 helices from Gi and Ras-AHD domains from Gαs was used to obtain well-performed purifications. All these formats of Gα subunits, including mini-DNGαs, DNGαs, DNGαi, and DNGα_si_ as well as human Gβ_1_, Gγ_2_, and a single-chain antibody scFv16^47,48^ were cloned into the pFastBac vector.

### Complexes expression and purification

For the D1R-G_s_ complex, the recombinant baculoviruses of DRD1, miniG_αs_, G_β1_ and G_γ2_ were prepared individually following the manufacturer’s instructions about the Bac-to-Bac baculovirus expression system (Thermo Fisher). Prior to protein expression, Sf9 cell cultures were grown to cell density at about 4×10^6^ cells/ mL in ESF 921 serum-free medium (Expression Systems). Subsequently, the Sf9 cells were co-infected with the four types of baculoviruses prepared above at the ratio of 1:1:1:1. After infection for 48h, the cultures were harvested and frozen at −80°C for further usage. Before purification of the D1R-G_s_ signaling complex, the stabilizing nanobody, Nb35, was prepared through the previously described method ^49^, fast-frozen by liquid nitrogen and stored at −80°C. For the purification of rotigotine-D1R-miniG_s_ complex, cell pellet of 1L culture was thawed at room temperature. The pellet was then resuspended in buffer containing 20 mM HEPES pH 7.3, 75 mM NaCl, 5 mM CaCl_2_, 5 mM MgCl_2_, 10% Glycerol, 0.3 mM TECP, protease inhibitor cocktail (Bimake, 1 mL/ 100 mL suspension). The protein complex was assembled on membrane by adding 100 μM rotigotine (TargetMol) and 10 μg/mL Nb35, which was added to stabilize the signaling complex. After incubation for half an hour, the suspension was treated with apyrase (25 mU mL^−1^, NEB) and incubated for another 1 hour at room temperature. The membrane in suspension was then solubilized by 0.5% Lauryl Maltose Neopentyl Glycol (L-MNG, Anatrace), 0.1% (w/v) cholesteryl hemisuccinate TRIS salt (CHS, Anatrace), 0.025% (w/v) digitonin (Biosynth). The membrane was solubilized for 3 hours at 4°C before separation by ultracentrifugation at 100,000 g (Ti45, Beckman) for 45 min. The isolated supernatant was incubated for 2 hours at 4°C with pre-equilibrated FLAG resin (Smart-Lifesciences). Detergents were directly exchanged upon FLAG resin by two washing steps in buffer containing 20 mM HEPES, pH 7.3, 100 mM NaCl, 0.3 mM TCEP, 20 μM rotigotine, and supplemented with different detergents: first 0.01% L-MNG, 0.002% CHS, 0.025% digitonin, then 0.015% L-MNG, 0.005% GDN, 0.004% CHS, 0.025% digitonin for 10 column volumes, each. The protein complex was then eluted in buffer containing 20 mM HEPES, pH 7.3, 100 mM NaCl, 0.3 mM TCEP, 20 μM rotigotine, 0.015% L-MNG, 0.005% GDN, 0.004% CHS, 0.025% digitonin, 200 μg/mL FLAG peptide. The eluted protein was concentrated to 0.5 mL by centrifugal filters with a 100 kDa molecular weight cut-off (ThermoFisher) and then loaded onto a Superdex 200 10/300 GL Increase column (GE Healthcare). The separation column was pre-equilibrated and ran in buffer containing 20 mM HEPES, pH 7.3, 100 mM NaCl, 0.3 mM TCEP, 20 μM rotigotine, 0.00075% L-MNG, 0.00025% GDN, 0.0002% CHS, 0.025% digitonin. Fractions of monomeric complex were collected and concentrated for electron microscopy experiments.

For the D2R-Gi, D3R-Gi, D4R-Gi complexes, the D2R/D3R/D4R, DNGα_i_/ DNGα_si_, Gβ_1_, Gγ_2_, and scFv16 were co-expressed in *Trichoplusia ni* (Hi5) insect cells using the Bac-to-Bac Baculovirus Expression System (Invitrogen). The D5R, DN_miniGα_s_, Gβ_1_, Gγ_2_ were also co-expressed in Hi5 insect cells. In addition, the D1R, DN_Gα_s_, Gβ_1_, Gγ_2_ were co-expressed in Sf9 insect cells. Cell cultures were grown in ESF 921 medium (Expression Systems) to a density of 3 × 10^6^ cell/mL and then infected with the different types of baculoviruses. Cell culture was collected by centrifugation 48 h post-infection and stored at −80°C until use.

For the purification of D2R-Gi, D3R-Gi, D4R-Gi, and D5R-Gs complex, cell pellets were lysed by homogenization in 20 mM HEPES pH 7.4, 20 mM KCl and 10 mM MgCl_2_ supplemented with Protease Inhibitor Cocktail (Bimake). The sample was centrifuged at 65,000 g for 30 min, then the membranes were re-suspended in 20 mM HEPES pH 7.4, 100 mM NaCl, 20 mM KCl, 10 mM MgCl_2_, 5 mM CaCl_2_, 25 mU/mL Apyrase (Sigma) and 10 μM Rotigotine. After incubation at RT for 1 h, the membranes were solubilized by addition of 0.5% (w/v) DDM (Anatrace) and 0.1% (w/v) Cholesteryl Hemisuccinate TRIS salt (CHS, Anatrace) for 2 h at 4°C. The supernatant was cleared by centrifugation and incubated with TALON (Clontech) resin overnight. After binding, the resin was washed with 20 column volumes of 20 mM HEPES pH 7.4, 100 mM NaCl, 2 mM MgCl_2_, 0.01% (w/v) Lauryl Maltose Neopentylglycol (LMNG, Anatrace), 0.002% (w/v) CHS, 25 mM imidazole and 10 μM Rotigotine. The complex was eluted with 5 column volumes of 20 mM HEPES pH 7.4, 100 mM NaCl, 2 mM MgCl_2_, 0.01% (w/v) LMNG, 0.002% (w/v) CHS, 250 mM imidazole and 10 μM Rotigotine. The protein was then concentrated and loaded onto a Superdex 200 Increase 10/300 column (GE Healthcare) pre-equilibrated with buffer containing 20 mM HEPES pH 7.4, 100 mM NaCl, 0.00075% (w/v) LMNG, 0.00025% (w/v) glyco-diosgenin (GDN, Anatrace), 0.0002% (w/v) CHS and 10 μM Rotigotine. The fractions for the monomeric complex were collected and concentrated for electron microscopy experiments.

### Cryo-EM grid preparation and data collection

For the preparation of cryo-EM grids, 3 μl of the purified complexes at 20 mg/ml for the D2R-Rotigotine-G_i_ complex, 17 mg/ml for the D3R-Rotigotine-G_i_ complex, 13 mg/ml for the D4R-Rotigotine-G_i_ complex, 20 mg/ml for the D5R-Rotigotine-G_s_ complex and 15 mg/ml for the D1R-Rotigotine-G_s_ complex were applied onto a glow-discharged holey carbon grid (Quantifoil R1.2/1.3). Grids were plunge-frozen in liquid ethane using Vitrobot Mark IV (Thermo Fischer Scientific). Frozen grids were transferred to liquid nitrogen and stored for data acquisition. For the D1R-Gs complex, D3R-Gi complex, D4R-Gi complex, and D5R-Gs complex, automatic data collection was performed on a Titan Krios equipped with a Gatan K3 direct electron detector in the Cryo-Electron Microscopy Research Center, Shanghai Institute of Materia Medica, Chinese Academy of Sciences (Shanghai, China). Cryo-EM imaging was performed micrographs were recorded in counting mode at a dose rate of about 8.0 e/Å^2^/s with a defocus ranging from −1.0 to −3.0 μm using the SerialEM software ^50^. The total exposure time was 8 s and 40 frames were recorded per micrograph. A total of 6301, 5156, 3562 and 4746 movies were collected for D1R-Gs complex, D3R-Gi complex, D4R-Gi complex, and D5R-Gs complex, respectively. For the D2R-Gi complex, automatic data collection was performed on a Titan Krios at 300 kV using Gatan K2 Summit detector in the Center of Cryo-Electron Microscopy, Zhejiang University (Hangzhou, China). Cryo-EM imaging was performed micrographs were recorded in counting mode at a dose rate of about 8.0 e/Å^2^/s with a defocus ranging from −1.0 to −3.0 μm using the SerialEM software ^50^. The total exposure time was 8 s and 40 frames were recorded per micrograph. A total of 5324 movies were collected for D2R-Gs complex.

### Image processing and map construction

Dose-fractionated image stacks were aligned using MotionCor2.1 ^51^. Contrast transfer function (CTF) parameters for each micrograph were estimated by Gctf ^52^. Cryo-EM data processing was performed using RELION-3.0-beta2 ^53^.

For the D1R-G_S_ complex, particle selections for 2D and 3D classifications were performed on a binned dataset with a pixel size of 1.60 Å. Automated particle picking yielded 2,685,434 particles that were subjected to reference-free 2D classification to discard poorly defined particles, producing 2,145,182 particles. After 6 rounds of 3D classification, a well-defined subset containing 448,516 particles was used to obtain the final map using a pixel size of 0.80 Å. Further refinement produced a final map with an indicated global resolution of 3.2 Å at a Fourier shell correlation (FSC) of 0.143.

For the D2R-G_i_ complex, particle selections for 2D and 3D classifications were performed on a binned dataset with a pixel size of 2.028 Å. Automated particle picking yielded 7,064,860 particles that were subjected to reference-free 2D classification to discard poorly defined particles. After 2 rounds of 3D classification, two well-defined subsets were selected. The selected subsets were subsequently subjected to 2 rounds of 3D classification with a mask on the receptor. One subset shows the high-quality receptor density was selected, producing 140,237 particles. The selected subset was subsequently subjected to 3D refinement, CTF refinement, and Bayesian polishing. The final refinement generated a map with an indicated global resolution of 3.0 Å at a Fourier shell correlation of 0.143.

For the D3R-G_i_ complex, particle selections for 2D and 3D classifications were performed on a binned dataset with a pixel size of 2.142 Å. Automated particle picking yielded 8,770,602 particles that were subjected to reference-free 2D classification to discard poorly defined particles. After 3 rounds of 3D classification, three well-defined subsets with 1,786,008 particles were selected and subsequently subjected to 3D refinement, CTF refinement, and Bayesian polishing. The final refinement generated a map with an indicated global resolution of 2.7 Å at a Fourier shell correlation of 0.143.

For the D4R-G_i_ complex, particle selections for 2D and 3D classifications were performed on a binned dataset with a pixel size of 2.16 Å. Automated particle picking yielded 4,333,829 particles that were subjected to reference-free 2D classification to discard poorly defined particles. After 3D classification, two well-defined subsets were selected and subsequently subjected to 3D classification with a mask on the receptor. Two subset shows the high-quality receptor density was selected, producing 471,638 particles. The selected subset was subsequently subjected to 3D refinement, CTF refinement, and Bayesian polishing. The final refinement generated a map with an indicated global resolution of 3.2 Å at a Fourier shell correlation of 0.143.

For the D5R-G_s_ complex, particle selections for 2D and 3D classifications were performed on a binned dataset with a pixel size of 2.142 Å. Automated particle picking yielded 7,900,346 particles that were subjected to reference-free 2D classification to discard poorly defined particles. After 2 rounds of 3D classification, one well-defined subset was selected and subsequently subjected to additional 4 rounds of 3D classification. four subsets show the high-quality receptor density was selected, producing 2,652,297 particles. The selected subset was subsequently subjected to 3D refinement, CTF refinement, and Bayesian polishing. The final refinement generated a map with an indicated global resolution of 3.1 Å at a Fourier shell correlation of 0.143. Local resolution was determined using the Bsoft ^54^ package with half maps as input maps.

### Model building and refinement

The structure of the D1R-Gs-Apomarphine complex (PDB: 7JVR) was used as the initial model for model rebuilding and refinement against the electron microscopy maps of D1R-Gs-Rotigotine and D5R-Gs-Rotigotine complexes. The structure of the D2R-Gi-Bromocriptine complex (PDB: 7JVQ) was used as the initial model for model rebuilding and refinement against the electron microscopy maps of D2R-Gi-Rotigotine complexes. The structure of the D3R-Gi-PD128907 complex (PDB: 7CMV) was used as the initial model for model rebuilding and refinement against the electron microscopy maps of the D3R-Gi-Rotigotine and the D4R-Gi-Rotigotine complexes. The model was docked into the electron microscopy density map using Chimera ^55^, followed by iterative manual adjustment and rebuilding in COOT ^56^ and ISOLDE ^57^. Real space and reciprocal space refinements were performed using Phenix programs ^58^. The model statistics was validated using MolProbity ^59^. Structural figures were prepared in Chimera, ChimeraX ^60^ and PyMOL (https://pymol.org/2/). The final refinement statistics are provided in Table S1.

### Radioligand binding assays

Binding assays were performed using membranes from HEK293T (ATCC CRL-11268) cells transiently expressing wild-type Dopamine receptors. For D1R and D5R Binding assays were set up in 96-well plates in standard binding buffer (50 mM HEPES, 50 mM NaCl, 5 mM MgCl2, 0.5 mM EDTA, pH 7.4). Saturation binding assays with 0.5–5 nM [3H]-SCH23390 (Perkin-Elmer) in standard binding buffer were performed to determine equilibrium dissociation constant (Kd) and Bmax, whereas 10 μM final concentration of Butaclamol was used to define nonspecific binding. For D2R, D3R, and D4R, binding assays were set up in 96-well plates in standard binding buffer (50 mM Tris, 0.1 mM EDTA, 10 mM MgCl2, 0.1% (w/v) BSA, pH 7.40). Saturation binding assays with 0.5-5 nM [3H]Methylspiperone (Perkin-Elmer) in standard binding buffer were performed to determine equilibrium dissociation constant (Kd) and Bmax, whereas 10 uM final concentration of Chlorpromazine was used to define nonspecific binding. All reactions were incubated for 2 hours at room temperature in the dark and terminated by rapid vacuum filtration onto chilled 0.3% PEI-soaked GF/A filters (Perkin-Elmer) followed by three quick washes with cold washing buffer (50 mM Tris HCl, pH 7.40). Radioactivity counts were determined using a Wallac Trilux MicroBeta counter (Perkin-Elmer). Results were analyzed using GraphPad Prism 8.4 (Graphpad Software Inc., San Diego, CA) using “One site -- Total and nonspecific binding”. Competition assays were performed similar to saturation binding assays except that various concentrations of competitor were premixed with [3H]-SCH23390 or [3H]Methylspiperone (Perkin-Elmer) near the pre-determined equilibrium dissociation constant (Kd) and then incubated for 2 h at room temperature in the dark with membranes from HEK293T (ATCC CRL-11268) cells transiently expressing wild-type receptors. Results were analyzed using GraphPad Prism 8.4 (Graphpad Software Inc., San Diego, CA) using “One site – Fit Ki”.

### Gs-mediated Gs-cAMP Accumulation Assay

For receptors D1R and D5R, GS-mediated GS-cAMP accumulation assays were performed with HEK293T (ATCC CRL-11268) cells transiently expressing human D1R or D5R wild-type and mutant along with the cAMP biosensor GloSensor-22F (Promega). Cells were seeded (20 000 cells/35 μL/well) into white 384 clear-bottom, tissue culture plates in DMEM containing 1% (v/v) dialyzed fetal bovine serum (FBS). Next day, 3x drug dilutions were diluted in HBSS, 20 mM N-(2-hydroxyethyl) piperazine-N’-ethanesulfonic acid (HEPES), 0.3% (w/v) bovine serum albumin (BSA), 0.03% (w/v) ascorbic acid, pH 7.4. Media was decanted from 384 well plates and 20 μL of drug buffer (HBSS, 20 mM HEPES, pH 7.4) containing GloSensor reagent was added per well and allowed to equilibrate for at least 15 min at room temperature. Cells were then treated with 10 μL per well of 3× drug using a FLIPR (Molecular Devices). After 15 min, Gs-cAMP accumulation was read on a TriLux Microbeta (PerkinElmer) plate counter. For receptors D2R, D3R, and D4R, GS-mediated GS-cAMP accumulation assays were performed as above except in the inhibition mode using a final concentration of 100nM of isoproterenol added to the cells 15 minutes prior to the addition of drug. Data were analyzed using the sigmoidal log(agonist) vs. dose response or sigmoidal log(inhibitor) vs. dose response function built into GraphPad Prism 8.4.

### PRESTO-Tango GPCRome Screening

Screening of the compounds in the PRESTO-Tango GPCRome was performed as previously described ^61^ with slight modifications. First, HTLA cells were plated in poly-L-lysine-coated 384-well white plates in DMEM containing 1% dialyzed FBS for 6 h. Next, the cells were transfected with 20 ng per well PRESTO-Tango receptor DNAs overnight. The cells were then treated with 10 μM Rotigotine without changing the medium and incubated for another 24 h. Each target was designed to have four wells for basal and four wells for sample. The remaining steps of the PRESTO-Tango protocol were followed. The results were plotted as fold change in the average basal signaling activity against individual receptors in GraphPad v.9.0. Selective receptors were repeated as a full dose–response assay to confirm activity.

---

**Fig. S1.**
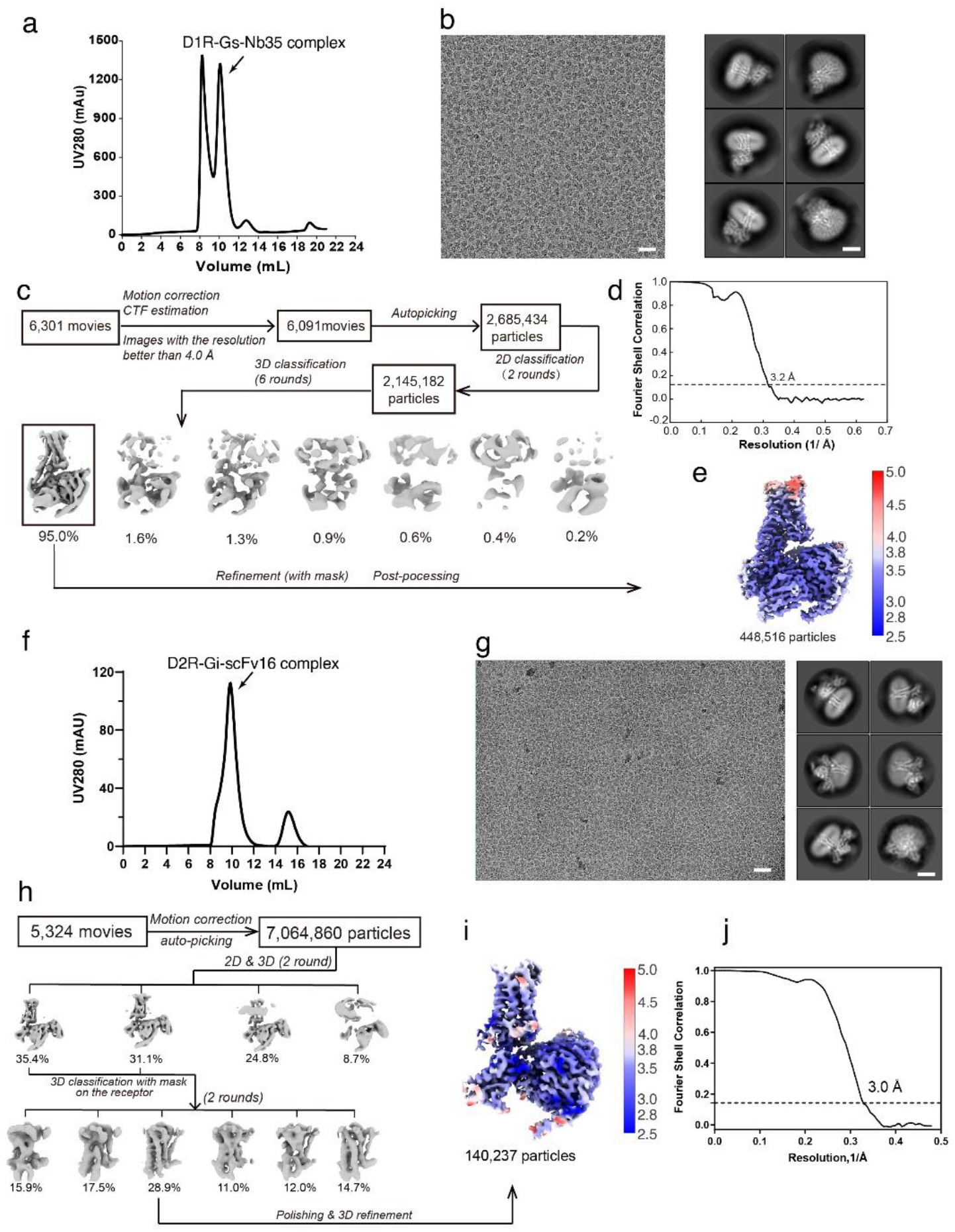
D1R-G_s_ and D2R-G_i_ complexes preparation and cryo-EM data processing. **a-e** D1R-G_s_ complexes preparation and cryo-EM data processing. Profiles of size-exclusion chromatography elution of the D1R-G_s_ complex (**a**). Representative cryo-EM images (scale bar, 50 nm) and 2D classification (scale bar, 5 nm) of the D1R-G_s_ complex (**b**). Workflow of cryo-EM single-particle analysis of the D1R-G_s_ complex (**c**). Fourier shell correlation curves of the D1R-G_s_ complex (**d**) Local resolution distribution of D1R-G_s_ complex (**e**). **f-j** D2R-G_i_ complexes preparation and cryo-EM data processing. Profiles of size-exclusion chromatography elution of the D2R-G_i_ complex (**f**). Representative cryo-EM images (scale bar, 50 nm) and 2D classification (scale bar, 5 nm) of the D2R-G_i_ complex (**g**). Workflow of cryo-EM single-particle analysis of the D2R-G_i_ complex (**h**). Local resolution distribution of D2R-G_i_ complex (**i**). Fourier shell correlation curves of the D2R-G_i_ complex (**j**).

**Fig. S2.**
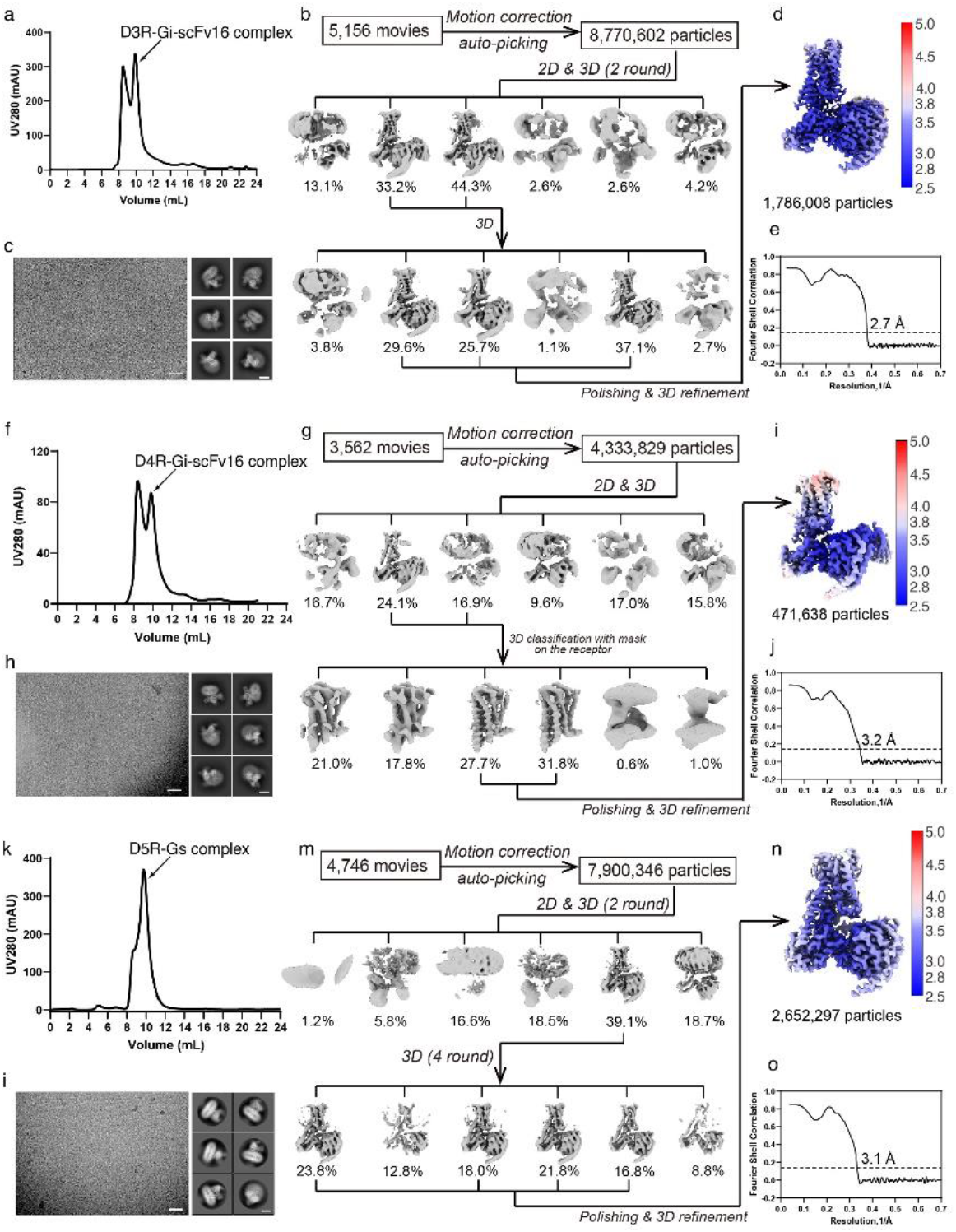
D3R-G_i_, D4R-G_i_, and D5R-G_s_ complexes preparation and cryo-EM data processing. **a-e** D3R-G_i_ complexes preparation and cryo-EM data processing. Profiles of size-exclusion chromatography elution of the D3R-G_i_ complex (**a**). Workflow of cryo-EM single-particle analysis of the D3R-G_i_ complex (**b**). Representative cryo-EM images (scale bar, 50 nm) and 2D classification (scale bar, 5 nm) of the D3R-G_i_ complex (**c**). Local resolution distribution of D3R-G_i_ complex (**d**). Fourier shell correlation curves of the D3R-G_i_ complex (**e**). **f-j** D4R-G_i_ complexes preparation and cryo-EM data processing. Profiles of size-exclusion chromatography elution of the D4R-G_i_ complex (**f**). Workflow of cryo-EM single-particle analysis of the D4R-G_i_ complex (**g**). Representative cryo-EM images (scale bar, 50 nm) and 2D classification (scale bar, 5 nm) of the D4R-G_i_ complex (**h**). Local resolution distribution of D4R-G_i_ complex (**i**). Fourier shell correlation curves of the D4R-G_i_ complex (**j**). **k-o** D5R-G_s_ complexes preparation and cryo-EM data processing. Profiles of size-exclusion chromatography elution of the D5R-G_s_ complex (**k**). Workflow of cryo-EM single-particle analysis of the D5R-G_s_ complex (**l**). Representative cryo-EM images (scale bar, 50 nm) and 2D classification (scale bar, 5 nm) of the D5R-G_s_ complex (**m**). Local resolution distribution of D5R-G_s_ complex (**n**). Fourier shell correlation curves of the D5R-G_s_ complex (**o**).

**Fig. S3.**
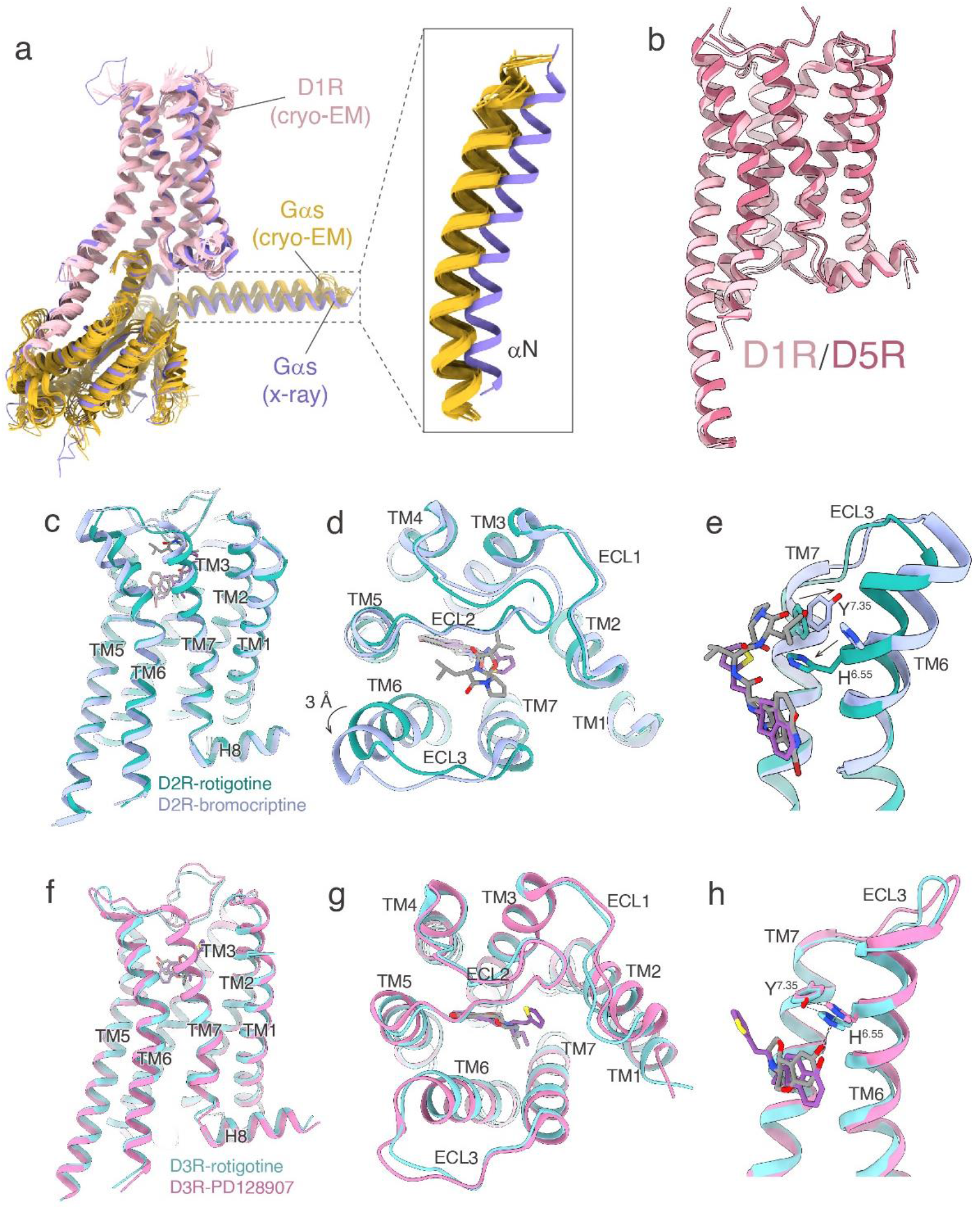
Structural comparison of active dopamine receptors. **a** Superposition of the cryo-EM D1R-Gs structures (pink) and the x-ay D1R-G_s_ structure (purple). **b** Superposition of D1R and D5R structure. **c-e** Structural features of active Rotigotine-D2R compared with that of bromocriptine-D2R (**a**). Comparison of ligand recognition between Rotigotine-D2R and bromocriptine-D2R (**b**). The structural difference of two active D2R in TM6 and TM7 (**c**). **f-h** Structural features of active Rotigotine-D3R compared with that of PD128907-D3R (**d**). Comparison of ligand recognition between Rotigotine-D3R and PD128907-D3R (**e**). The structural difference of two active D3R in TM6 and TM7 (**f**).

**Fig. S4.**
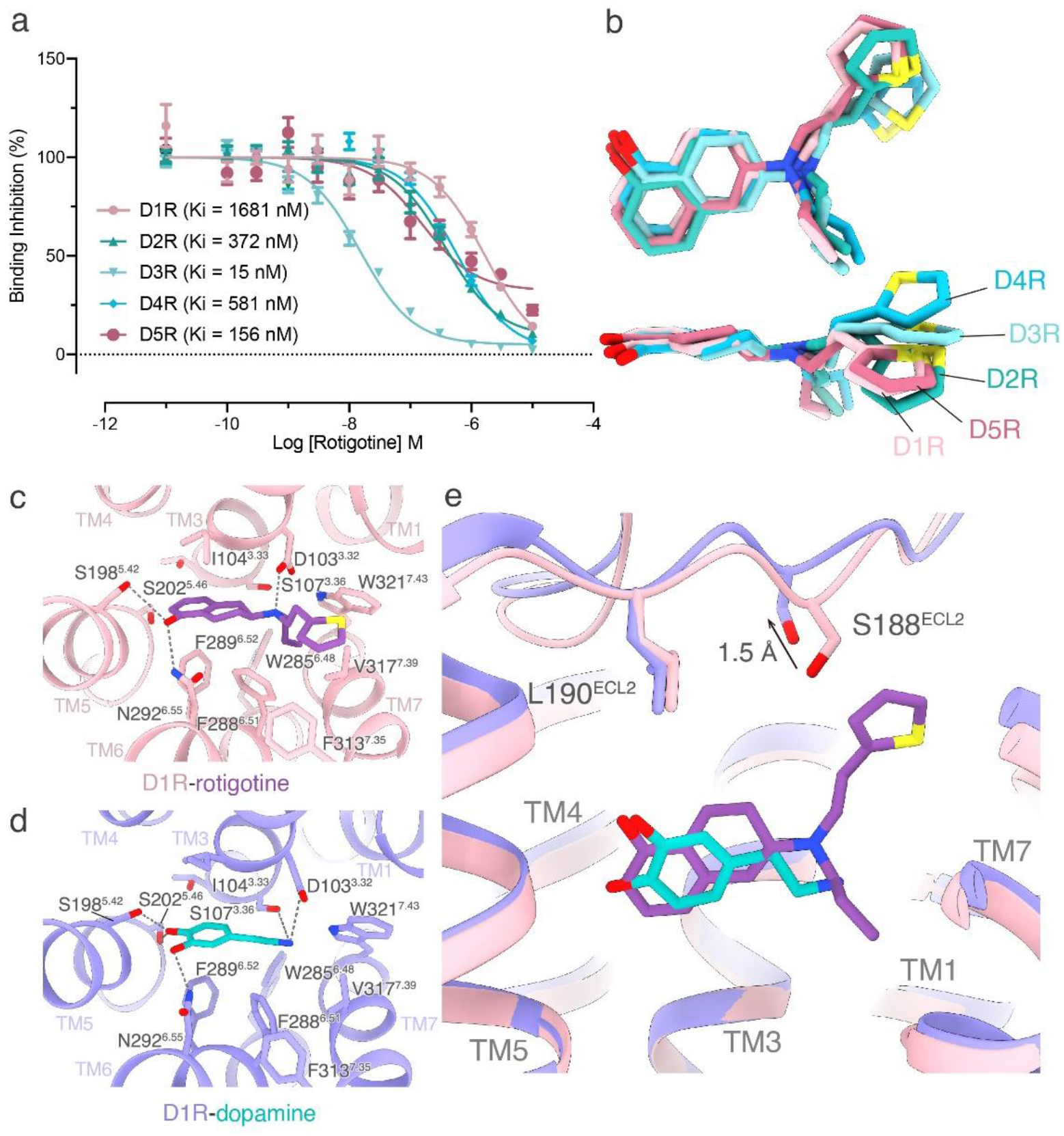
Comparison of Rotigotine conformation from five dopamine receptor and the comparison of the binding modes between Rotigotine and dopamine. **a** Rotigotine competition binding curve to WT Dopamine Receptors. Graphical representation of Rotigotine competition curves for wild-type receptors. Dose response curves were analyzed using “One site – Fit LogIC50” function in Graphpad Prism 8.4 software (Graphpad Software Inc., San Diego, CA). All data are presented as mean values ± standard error of measurement (SEM) with a minimum of four technical replicates and N = 3 biological replicates. **b** Superposition of Rotigotine structure from the five dopamine receptors. **c** The binding mode of Rotigotine in D1R. **d** The binding mode of dopamine in D1R. **e** Structural comparison of Rotigotine and dopamine in D1R.

**Fig. S5.**
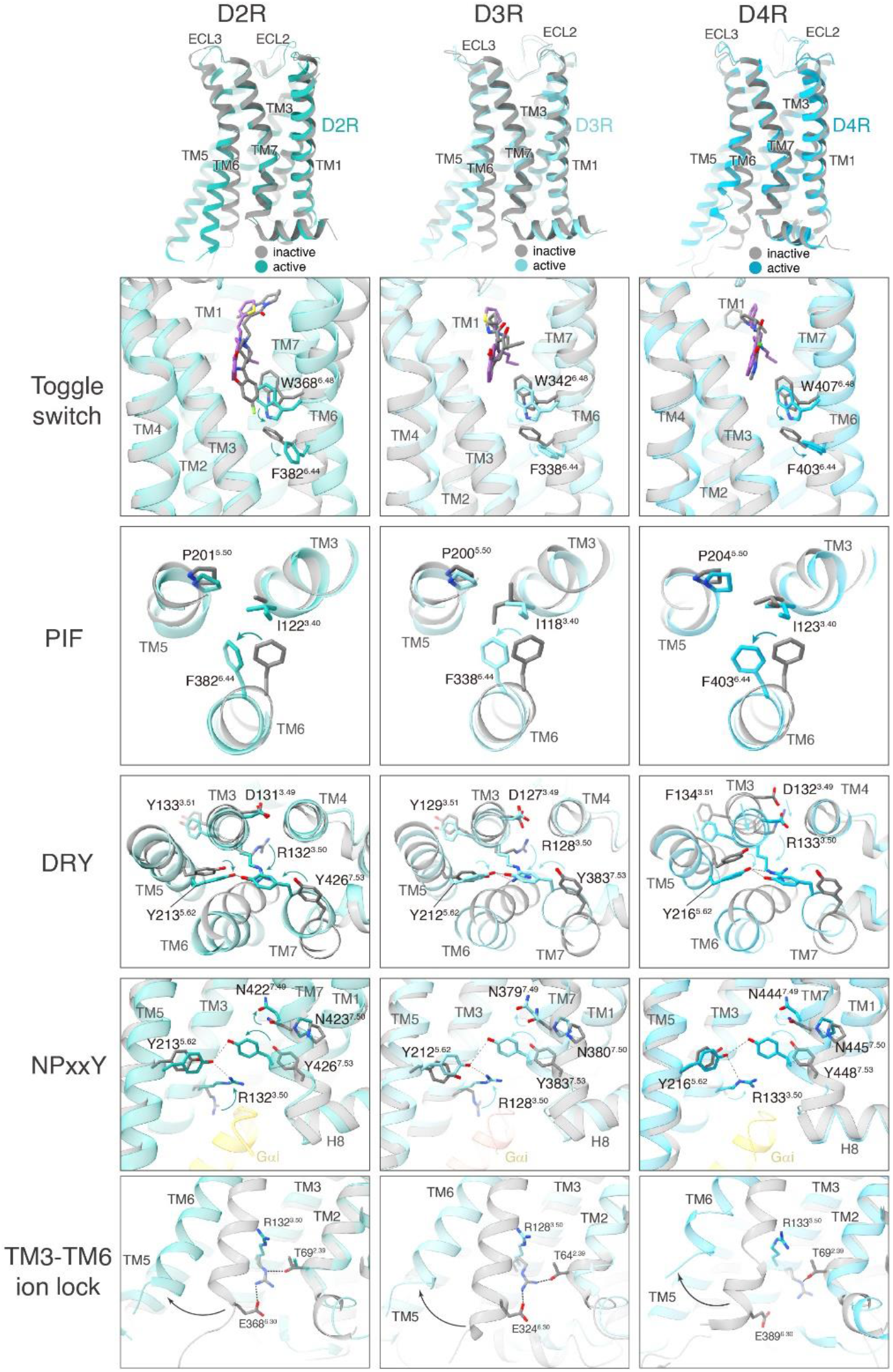
D2-like receptors activation. **a** Toggle switch induced by agonist binding in D2R, D3R, and D4R. **b** Conformational change of PIF motif in D2R, D3R, and D4R. c Conformational change of DRY motif in D2R, D3R, and D4R. **d** Rotation of the NPxxY motif in D2R, D3R, and D4R. **e** The breaking of TM3-TM6 ion lock of D2R, D3R, and D4R.

**Fig. S6.**
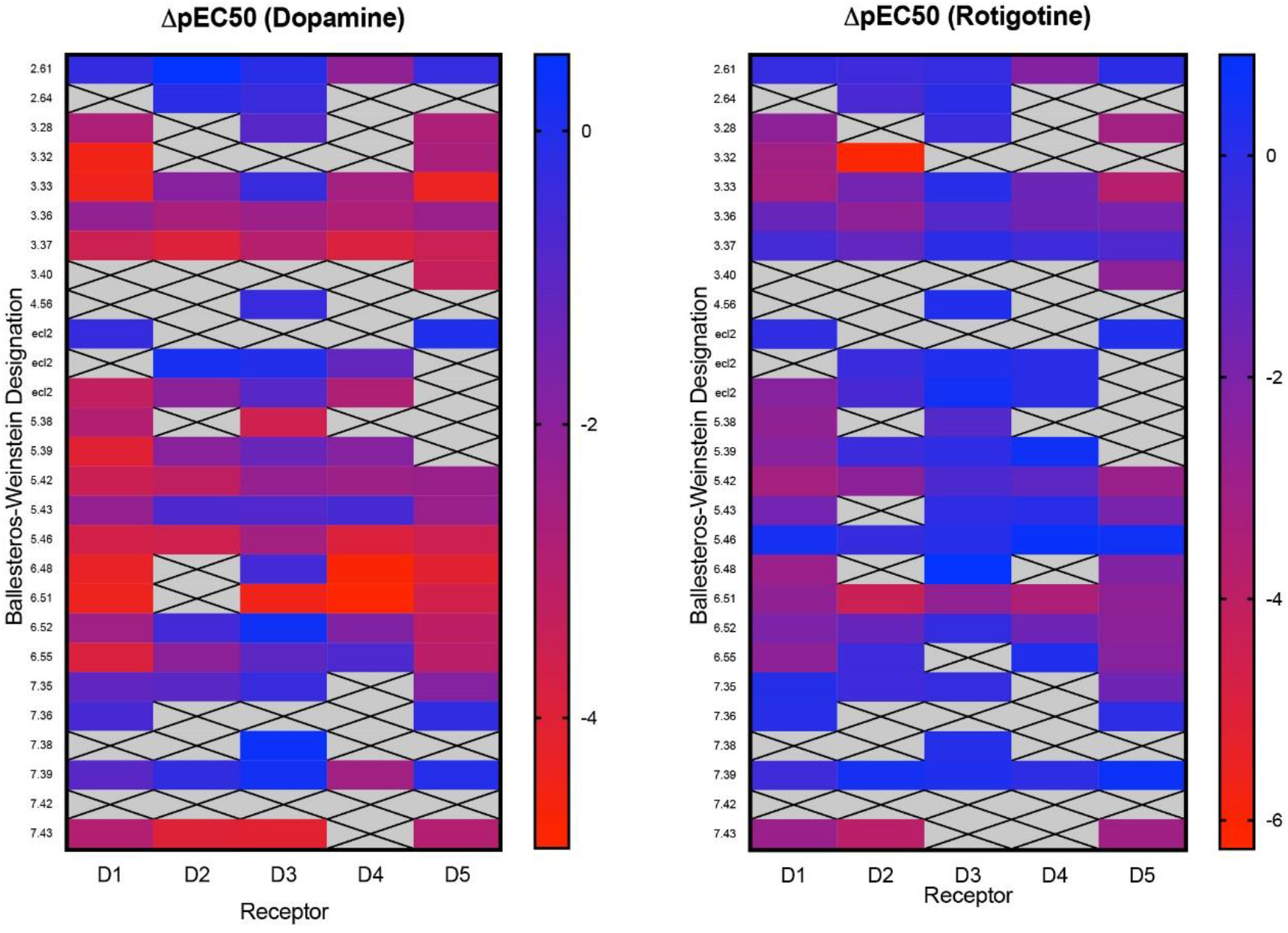
Heatmap for ligand changes in ΔpEC_50_ in response to stimulation. Ligand-specific interactions with dopamine receptors. Heatmap summary of mutagenesis studies showing the effects of orthosteric-site residues on functional activity with regards to dopamine and Rotigotine. Dose response curves were analyzed using “One site – Fit LogIC50” function in Graphpad Prism 8.4 software (Graphpad Software Inc., San Diego, CA). All data are presented as mean values ± standard error of measurement (SEM) with a minimum of four technical replicates and N = 3 biological replicates.

**Fig. S7.**
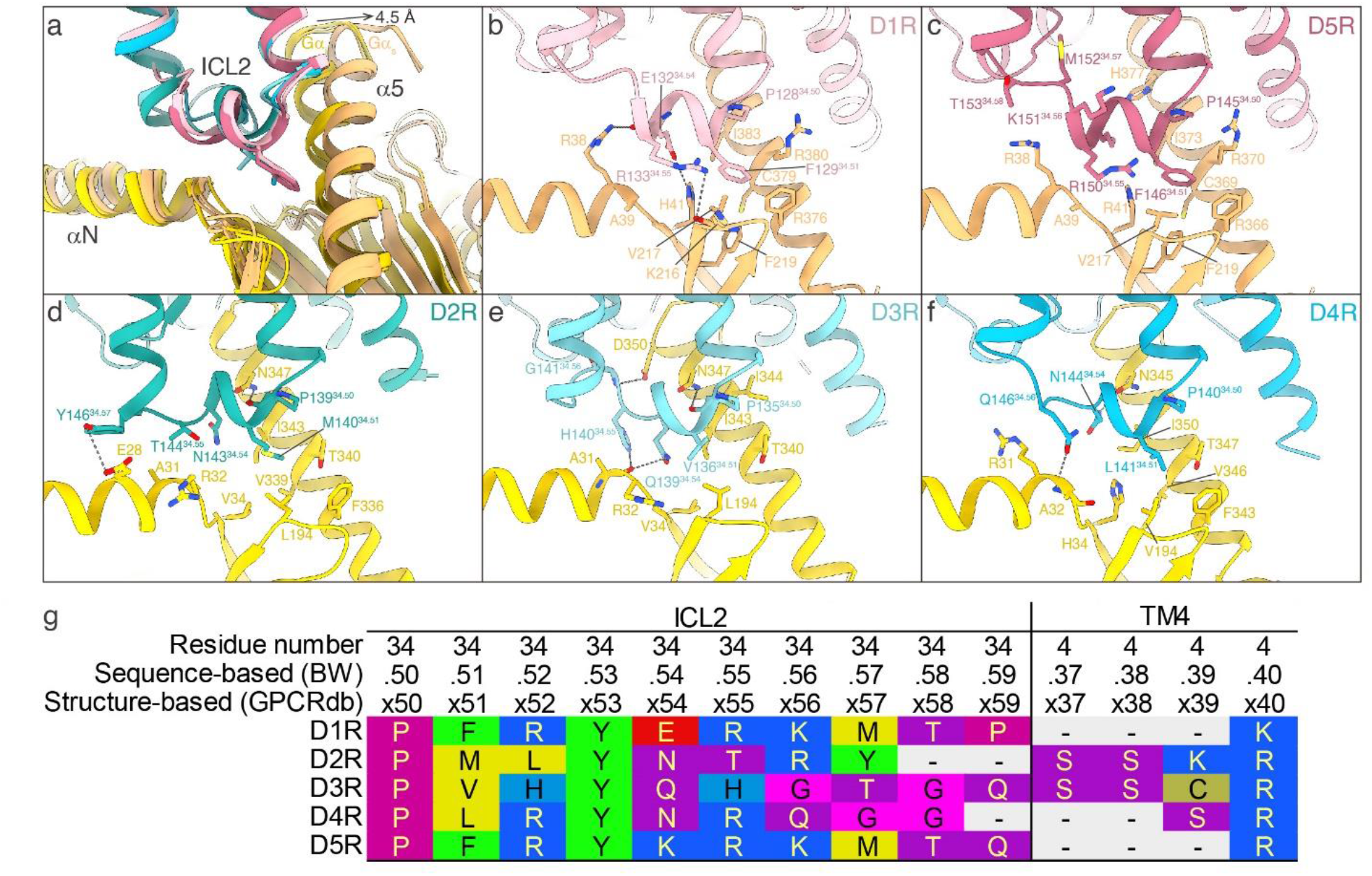
ICL2-G protein interaction of dopamine receptors. **a** Super position of the ICL2-G protein structure of the five dopamine receptor-G protein complexes. **b** D1R ICL2-G protein interaction. **c** D2R ICL2-G protein interaction. **d** D3R ICL2-G protein interaction. **e** D4R ICL2-G protein interaction. **f** D5R ICL2-G protein interaction. **g** ICL2 sequence alignment of dopamine receptors.

**Fig. S8.**
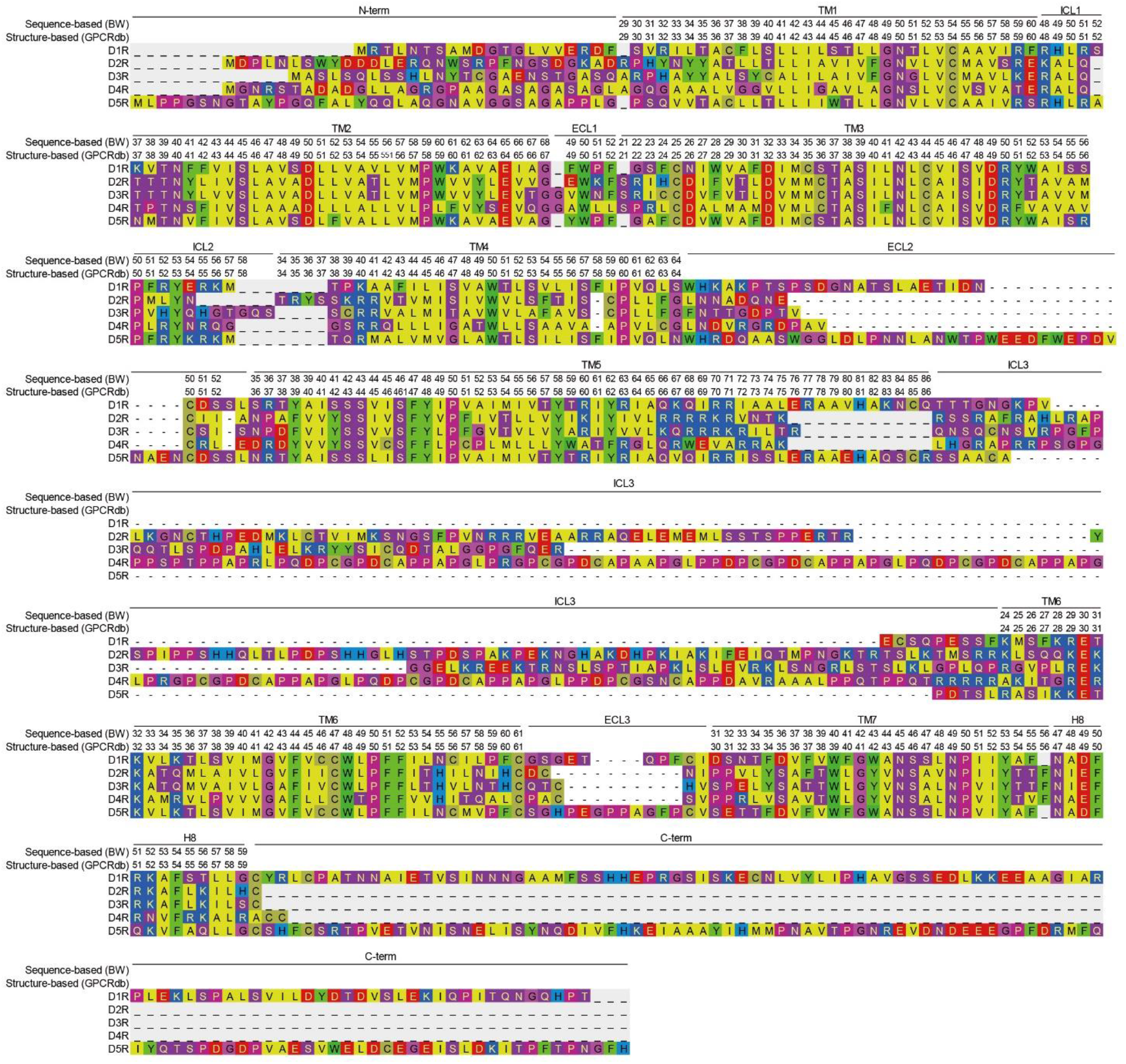
Sequence alignment of dopamine receptors (gpcrdb.org)^62^.

**Table S1.**
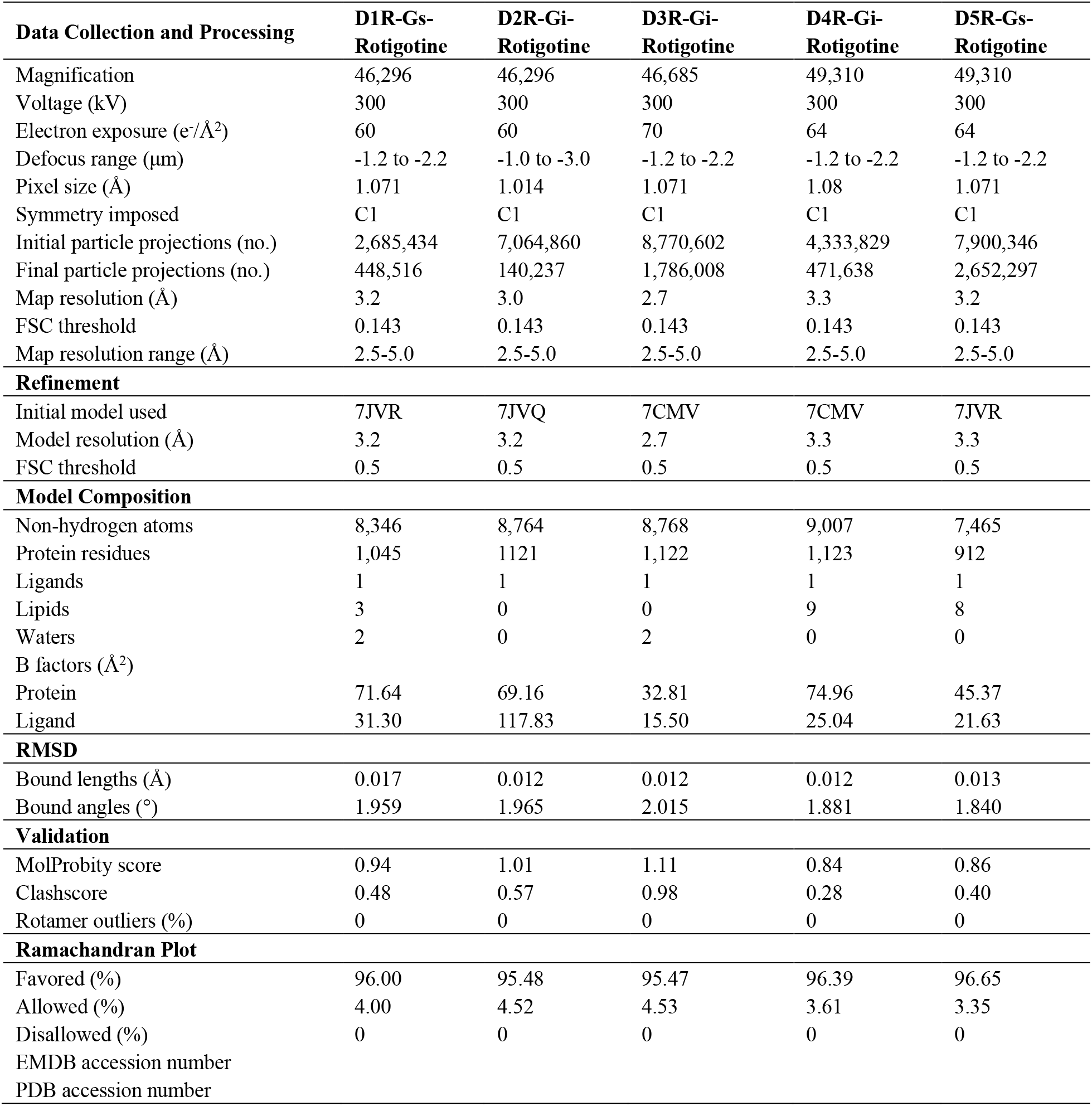
Cryo-EM data collection, model refinement, and validation statistics.

| **Data Collection and Processing** | **D1R-Gs-Rotigotine** | **D2R-Gi-Rotigotine** | **D3R-Gi-Rotigotine** | **D4R-Gi-Rotigotine** | **D5R-Gs-Rotigotine** |
| --- | --- | --- | --- | --- | --- |
| Magnification | 46,296 | 46,296 | 46,685 | 49,310 | 49,310 |
| Voltage (kV) | 300 | 300 | 300 | 300 | 300 |
| Electron exposure (e<sup>-</sup>/Å<sup>2</sup>) | 60 | 60 | 70 | 64 | 64 |
| Defocus range (μm) | -1.2 to -2.2 | -1.0 to -3.0 | -1.2 to -2.2 | -1.2 to -2.2 | -1.2 to -2.2 |
| Pixel size (Å) | 1.071 | 1.014 | 1.071 | 1.08 | 1.071 |
| Symmetry imposed | C1 | C1 | C1 | C1 | C1 |
| Initial particle projections (no.) | 2,685,434 | 7,064,860 | 8,770,602 | 4,333,829 | 7,900,346 |
| Final particle projections (no.) | 448,516 | 140,237 | 1,786,008 | 471,638 | 2,652,297 |
| Map resolution (Å) | 3.2 | 3.0 | 2.7 | 3.3 | 3.2 |
| FSC threshold | 0.143 | 0.143 | 0.143 | 0.143 | 0.143 |
| Map resolution range (Å) | 2.5-5.0 | 2.5-5.0 | 2.5-5.0 | 2.5-5.0 | 2.5-5.0 |
| **Refinement** |  |  |  |  |  |
| Initial model used | 7JVR | 7JVQ | 7CMV | 7CMV | 7JVR |
| Model resolution (Å) | 3.2 | 3.2 | 2.7 | 3.3 | 3.3 |
| FSC threshold | 0.5 | 0.5 | 0.5 | 0.5 | 0.5 |
| **Model Composition** |  |  |  |  |  |
| Non-hydrogen atoms | 8,346 | 8,764 | 8,768 | 9,007 | 7,465 |
| Protein residues | 1,045 | 1121 | 1,122 | 1,123 | 912 |
| Ligands | 1 | 1 | 1 | 1 | 1 |
| Lipids | 3 | 0 | 0 | 9 | 8 |
| Waters | 2 | 0 | 2 | 0 | 0 |
| B factors (Å<sup>2</sup>) |  |  |  |  |  |
| Protein | 71.64 | 69.16 | 32.81 | 74.96 | 45.37 |
| Ligand | 31.30 | 117.83 | 15.50 | 25.04 | 21.63 |
| **RMSD** |  |  |  |  |  |
| Bound lengths (Å) | 0.017 | 0.012 | 0.012 | 0.012 | 0.013 |
| Bound angles (°) | 1.959 | 1.965 | 2.015 | 1.881 | 1.840 |
| **Validation** |  |  |  |  |  |
| MolProbity score | 0.94 | 1.01 | 1.11 | 0.84 | 0.86 |
| Clashscore | 0.48 | 0.57 | 0.98 | 0.28 | 0.40 |
| Rotamer outliers (%) | 0 | 0 | 0 | 0 | 0 |
| **Ramachandran Plot** |  |  |  |  |  |
| Favored (%) | 96.00 | 95.48 | 95.47 | 96.39 | 96.65 |
| Allowed (%) | 4.00 | 4.52 | 4.53 | 3.61 | 3.35 |
| Disallowed (%) | 0 | 0 | 0 | 0 | 0 |
| EMDB accession number |  |  |  |  |  |
| PDB accession number |  |  |  |  |  |

**Table S2.**
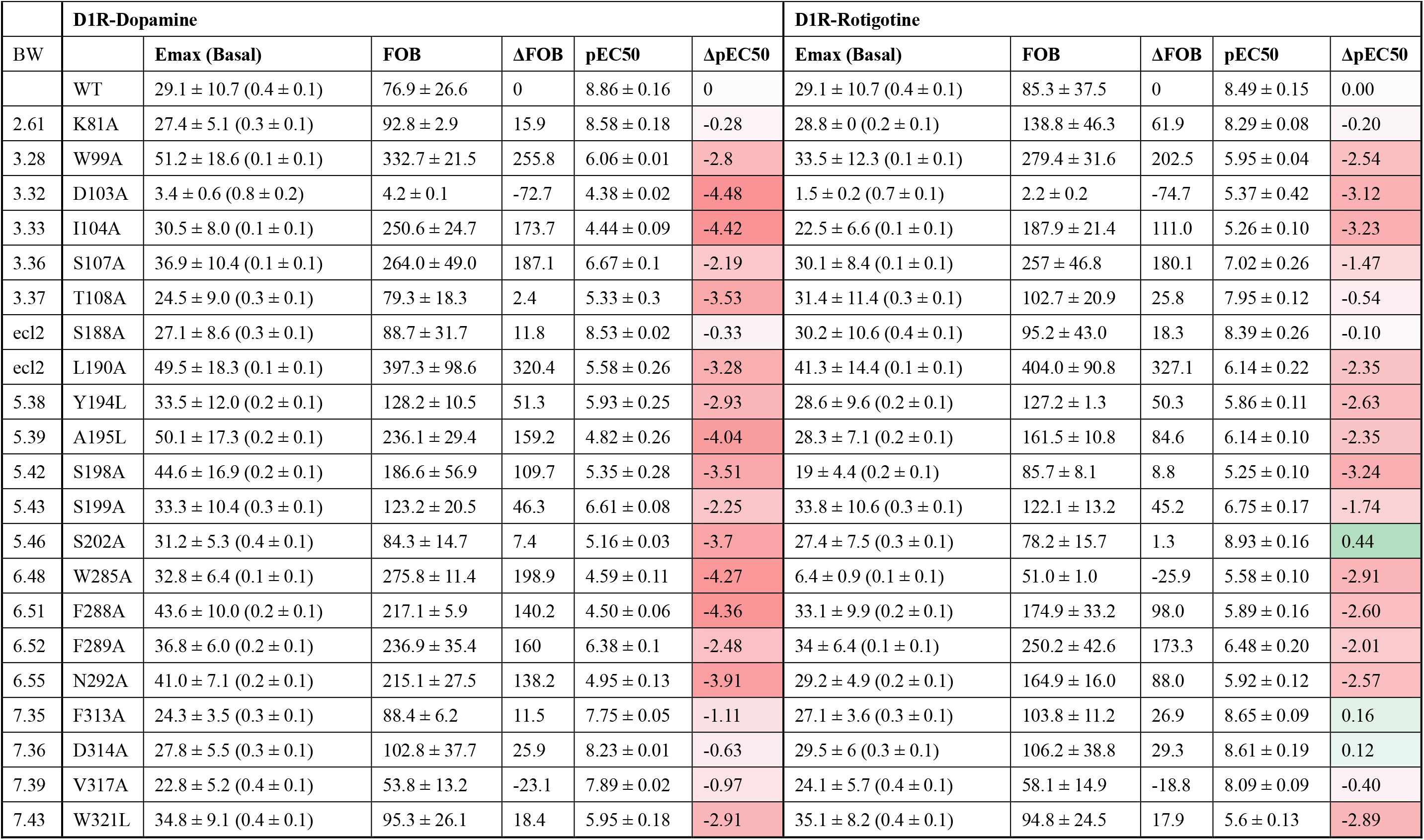
Gs-mediated cAMP accumulation assay results of D1R. Potency (pEC50) were extracted from a minimum of 3 independent assays in at least triplicate. pEC50 displayed values are mean ± SEM. Delta FOB for difference in either Fold of Basal (FOB) or Delta pEC50 when compared to wild-type receptor value. Average Emax and basal values were determined from “log(agonist) vs. response – Variable slope (four parameters) or log(inhibition) vs. response – Variable slope (four parameters)” function in Graphpad Prism 8.4 software (Graphpad Software Inc., San Diego, CA) and were divided by 10^3^ for display, basal values are enclosed with parentheses in the table column. Color scheme is based on the effects of mutations on relative pEC50 and Fold of Basal (FOB) values with red for reduced potency/efficacy and blue for increased potency/efficacy when compared to wild-type values for each ligand. ND - not determined.

**Table S3.** Gi-mediated cAMP accumulation assay results of D2R. Potency (pEC50) were extracted from a minimum of 3 independent assays in at least triplicate. pEC50 displayed values are mean ± SEM. Delta FOB for difference in either Fold of Basal (FOB) or Delta pEC50 when compared to wild-type receptor value. Average Emax and basal values were determined from “log(agonist) vs. response – Variable slope (four parameters) or log(inhibition) vs. response – Variable slope (four parameters)” function in Graphpad Prism 8.4 software (Graphpad Software Inc., San Diego, CA) and were divided by 10^3^ for display, basal values are enclosed with parentheses in the table column. Color scheme is based on the effects of mutations on relative pEC50 and Fold of Basal (FOB) values with red for reduced potency/efficacy and blue for increased potency/efficacy when compared to wild-type values for each ligand. ND - not determined.

|  | D2-Dopamine |  |  |  |  |  | D2R-Rotigotine |  |  |  |  |
| --- | --- | --- | --- | --- | --- | --- | --- | --- | --- | --- | --- |
| BW |  | Emax (Basal) | FOB | ΔFOB | pEC50 | ΔpEC50 | Emax (Basal) | FOB | ΔFOB | pEC50 | ΔpEC50 |
|  | WT | 1.3 ± 0.5 (10 ± 3.8) | 7.1 ± 0.5 | 0 | 9.59 ± 0.11 | 0 | 1.8 ± 0.4 (13.8 ± 4) | 7.6 ± 0.9 | 0 | 11.1 ± 0.15 | 0.00 |
| 2.61 | V91A | 2.1 ± 0.9 (13 ± 5.1) | 6.3 ± 0.2 | -0.8 | 10.11 ± 0.06 | 0.52 | 2.7 ± 0.1 (18 ± 0.1) | 6.5 ± 0.7 | -0.6 | 10.73 ± 0.15 | -0.37 |
| 2.64 | L94A | 1.9 ± 0.9 (10.3 ± 5.1) | 5 ± 0.3 | -2.1 | 9.47 ± 0.06 | -0.12 | 2.7 ± 0.7 (14 ± 4.5) | 4.8 ± 0.6 | -2.3 | 10.45 ± 0.17 | -0.65 |
| 3.32 | D114A | N.D. | N.D. | N.D. | N.D. | N.D. | 5.8 ± 2.0 (18.2 ± 6.1) | 3.1 ± 0.1 | -4.0 | 4.84 ± 0.22 | -6.26 |
| 3.33 | V115A | 2.2 ± 0.6 (13.4 ± 4.7) | 5.8 ± 0.4 | -1.3 | 7.65 ± 0.02 | -1.94 | 2.7 ± 0.6 (15.8 ± 3.5) | 5.8 ± 0.3 | -1.3 | 9.35 ± 0.06 | -1.75 |
| 3.36 | C118A | 6.0 ± 1.4 (19.1 ± 4.1) | 3.2 ± 0.1 | -3.9 | 6.9 ± 0.36 | -2.69 | 4.4 ± 0.4 (22.2 ± 4.3) | 4.9 ± 0.6 | -2.2 | 8.55 ± 0.02 | -2.55 |
| 3.37 | T119A | 4.9 ± 0.8 (11.8 ± 3.4) | 2.3 ± 0.3 | -4.8 | 5.64 ± 0.07 | -3.95 | 1.8 ± 0.3 (13.7 ± 3.0) | 7.2 ± 0.5 | 0.1 | 9.79 ± 0.19 | -1.31 |
| ecl2 | I183A | 2.1 ± 0.4 (12.8 ± 3.5) | 5.7 ± 0.6 | -1.4 | 9.74 ± 0.01 | 0.15 | 2.1 ± 0.3 (14.0 ± 3.2) | 6.4 ± 0.7 | -0.7 | 10.82 ± 0.20 | -0.28 |
| ecl2 | I184A | 2.3 ± 0.5 (9.3 ± 3) | 3.8 ± 0.4 | -3.3 | 7.58 ± 0.07 | -2.01 | 2.1 ± 0.4 (11.6 ± 3.0) | 5.1 ± 0.7 | -2.0 | 10.49 ± 0.19 | -0.61 |
| 5.39 | V190A | 1.7 ± 0.6 (9.7 ± 3.8) | 5.4 ± 0.3 | -1.7 | 7.61 ± 0.08 | -1.98 | 2.1 ± 0.5 (12.8 ± 3.9) | 5.7 ± 0.6 | -1.4 | 10.75 ± 0.22 | -0.35 |
| 5.42 | S193A | 2.4 ± 0.3 (14 ± 4.6) | 5.5 ± 1.5 | -1.6 | 6.34 ± 0.03 | -3.25 | 2.4 ± 0.4 (15.7 ± 3.6) | 6.1 ± 0.6 | -1.0 | 8.60 ± 0.09 | -2.50 |
| 5.43 | S194A | 2.1 ± 0.3 (14.5 ± 4.8) | 6.3 ± 1.2 | -0.8 | 8.82 ± 0.11 | -0.77 | N.D. | N.D. | N.D. | N.D. | N.D. |
| 5.46 | S197A | 7.5 ± 2.8 (12.3 ± 3.7) | 1.8 ± 0.2 | -5.3 | 6 ± 0.02 | -3.59 | 1.8 ± 0.4 (12.6 ± 3.9) | 6.8 ± 0.7 | -0.3 | 10.88 ± 0.15 | -0.22 |
| 6.48 | W380A | N.D. | N.D. | N.D. | N.D. | N.D. | N.D. | N.D. | N.D. | N.D. | N.D. |
| 6.51 | F389A | N.D. | N.D. | N.D. | N.D. | N.D. | 3.4 ± 1.0 (9.9 ± 3.5) | 2.7 ± 0.4 | -4.4 | 6.67 ± 0.43 | -4.43 |
| 6.52 | F390A | 1.8 ± 0.4 (10.9 ± 2.7) | 6.2 ± 0.1 | -0.9 | 9.03 ± 0.26 | -0.56 | 2.0 ± 0.5 (11.5 ± 2.7) | 5.9 ± 0.2 | -1.2 | 9.72 ± 0.13 | -1.38 |
| 6.55 | H393A | 2.8 ± 0.7 (15.1 ± 4.4) | 5.3 ± 0.2 | -1.8 | 7.56 ± 0.16 | -2.03 | 2.3 ± 0.4 (14.2 ± 3.8) | 6.0 ± 0.8 | -1.1 | 10.74 ± 0.13 | -0.36 |
| 7.35 | Y408A | 1.3 ± 0.3 (8.6 ± 1.9) | 6.4 ± 0.2 | -0.7 | 8.6 ± 0.12 | -0.99 | 1.5 ± 0.3 (9.2 ± 1.9) | 6.1 ± 0.6 | -1.0 | 10.72 ± 0.08 | -0.38 |
| 7.39 | T412A | 1.6 ± 0.5 (12.8 ± 4.6) | 8.4 ± 1.2 | 1.3 | 9.33 ± 0.16 | -0.26 | 2.3 ± 0.6 (14.4 ± 4.5) | 6.0 ± 0.5 | -1.1 | 11.57 ± 0.13 | 0.47 |
| 7.43 | Y416A | 3.3 ± 0.5 (11.9 ± 3.5) | 3.4 ± 0.6 | -3.7 | 5.59 ± 0.12 | -4 | 11.2 ± 0.6 (17 ± 0.3) | 1.5 ± 0.1 | -5.6 | 7.23 ± 0.14 | -3.87 |

**Table S4.** Gi-mediated cAMP accumulation assay results of D3R. Potency (pEC50) were extracted from a minimum of 3 independent assays in at least triplicate. pEC50 displayed values are mean ± SEM. Delta FOB for difference in either Fold of Basal (FOB) or Delta pEC50 when compared to wild-type receptor value. Average Emax and basal values were determined from “log(agonist) vs. response – Variable slope (four parameters) or log(inhibition) vs. response – Variable slope (four parameters)” function in Graphpad Prism 8.4 software (Graphpad Software Inc., San Diego, CA) and were divided by 10^3^ for display, basal values are enclosed with parentheses in the table column. Color scheme is based on the effects of mutations on relative pEC50 and Fold of Basal (FOB) values with red for reduced potency/efficacy and blue for increased potency/efficacy when compared to wild-type values for each ligand. ND - not determined.

|  | D3-Dopamine |  |  |  |  |  | D3R-Rotigotine |  |  |  |  |
| --- | --- | --- | --- | --- | --- | --- | --- | --- | --- | --- | --- |
| BW |  | Emax (Basal) | FOB | ΔFOB | pEC50 | ΔpEC50 | Emax (Basal) | FOB | ΔFOB | pEC50 | ΔpEC50 |
|  | WT | 2.0 ± 0.1 (8.4 ± 0.2) | 4.2 ± 0.1 | 0 | 9.94 ± 0.12 | 0 | 1.8 ± 0.2 (6.2 ± 0.5) | 3.5 ± 0.1 | 0 | 9.56 ± 0.02 | 0.00 |
| 2.61 | V86A | 2.1 ± 0.2 (7.3 ± 0.6) | 3.4 ± 0.1 | -0.8 | 9.83 ± 0.05 | -0.11 | 2.4 ± 0.1 (6.6 ± 0.1) | 2.8 ± 0.1 | -1.4 | 9.38 ± 0.06 | -0.18 |
| 2.64 | L89A | 2.0 ± 0.3 (3.2 ± 0.6) | 1.6 ± 0.1 | -2.6 | 9.55 ± 0.46 | -0.39 | 2.7 ± 0.5 (3.5 ± 0.7) | 1.3 ± 0.1 | -2.9 | 9.61 ± 0.13 | 0.05 |
| 3.28 | F106A | 4.5 ± 0.8 (8.5 ± 1.5) | 1.9 ± 0 | -2.3 | 9.03 ± 0.41 | -0.91 | 3.7 ± 0.7 (5.6 ± 0.9) | 1.6 ± 0.1 | -2.6 | 9.21 ± 0.13 | -0.35 |
| 3.32 | D110A | N.D. | N.D. | N.D. | N.D. | N.D. | N.D. | N.D. | N.D. | N.D. | N.D. |
| 3.33 | V111A | 2.3 ± 0.1 (8.3 ± 0.1) | 3.6 ± 0.2 | -0.6 | 9.57 ± 0.06 | -0.37 | 2.2 ± 0.1 (6.7 ± 0.3) | 3.1 ± 0.2 | -1.1 | 9.67 ± 0.07 | 0.11 |
| 3.36 | C114A | 1.4 ± 0.2 (5.8 ± 0.7) | 4.1 ± 0.2 | -0.1 | 7.54 ± 0.07 | -2.4 | 1.6 ± 0.2 (3.8 ± 0.5) | 2.3 ± 0.1 | -1.9 | 8.63 ± 0.02 | -0.93 |
| 3.37 | T115A | 3.9 ± 0.3 (6.9 ± 0.6) | 1.8 ± 0.1 | -2.4 | 6.93 ± 0.12 | -3.01 | 2.9 ± 0.4 (6.6 ± 0.8) | 2.3 ± 0.1 | -1.9 | 9.62 ± 0.06 | 0.06 |
| 4.56 | V164A | 1.7 ± 0.6 (4.6 ± 1.5) | 2.6 ± 0.1 | -1.6 | 9.57 ± 0.15 | -0.37 | 2.4 ± 0.6 (5.5 ± 1.5) | 2.3 ± 0.1 | -1.9 | 9.81 ± 0.02 | 0.25 |
| ecl2 | C181A | N.D. | N.D. | N.D. | N.D. | N.D. | N.D. | N.D. | N.D. | N.D. | N.D. |
| ecl2 | S182A | 1.7 ± 0.5 (6.1 ± 1.7) | 3.6 ± 0.1 | -0.6 | 9.91 ± 0.09 | -0.03 | 2.0 ± 0.5 (6.6 ± 1.6) | 3.3 ± 0.2 | -0.9 | 9.85 ± 0.03 | 0.29 |
| ecl2 | I183A | 2.2 ± 0.1 (8.1 ± 1.0) | 3.6 ± 0.2 | -0.6 | 9 ± 0.12 | -0.94 | 2.2 ± 0.3 (8.0 ± 1.2) | 3.5 ± 0.2 | -0.7 | 10.09 ± 0.04 | 0.53 |
| 5.38 | F188A | 2.3 ± 0.1 (8.3 ± 0.3) | 3.6 ± 0.1 | -0.6 | 6.34 ± 0.03 | -3.6 | 2.5 ± 0.1 (5.6 ± 0.2) | 2.2 ± 0.1 | -2.0 | 8.62 ± 0.06 | -0.94 |
| 5.39 | V189A | 3.2 ± 0.5 (8.7 ± 1.4) | 2.7 ± 0.1 | -1.5 | 8.64 ± 0.05 | -1.3 | 3.0 ± 0.5 (8.2 ± 1.7) | 2.6 ± 0.2 | -1.6 | 9.53 ± 0.04 | -0.03 |
| 5.42 | S192A | 2.7 ± 0.5 (7.5 ± 1.8) | 2.8 ± 0.2 | -1.4 | 7.7 ± 0.09 | -2.24 | 3.3 ± 0.6 (7.4 ± 1.7) | 2.2 ± 0.2 | -2.0 | 8.82 ± 0.07 | -0.74 |
| 5.43 | S193A | 1.6 ± 0.2 (3.9 ± 0.8) | 2.4 ± 0.3 | -1.8 | 9.11 ± 0.46 | -0.83 | 1.5 ± 0.2 (3.8 ± 0.8) | 2.4 ± 0.2 | -1.8 | 9.53 ± 0.10 | -0.03 |
| 5.46 | S196A | 11.4 ± 1.5 (14.1 ± 1.8) | 1.2 ± 0 | -3 | 7.37 ± 0.19 | -2.57 | 4.0 ± 0.5 (12.9 ± 2.5) | 3.2 ± 0.3 | -1.0 | 9.69 ± 0.04 | 0.13 |
| 6.48 | W342A | 5.5 ± 0.5 (6.8 ± 0.8) | 1.2 ± 0 | -3 | 9.38 ± 0.47 | -0.56 | 4.6 ± 0.7 (5.6 ± 0.8) | 1.2 ± 0.1 | -3.0 | 10.47 ± 0.35 | 0.91 |
| 6.51 | F345A | 2.0 ± 0.3 (8.1 ± 0.8) | 4.2 ± 0.5 | 0 | 5.49 ± 0.02 | -4.45 | 2.6 ± 0.3 (6.4 ± 0.8) | 2.5 ± 0.1 | -1.7 | 6.91 ± 0.04 | -2.65 |
| 6.52 | F346A | 2.5 ± 0.2 (7.6 ± 0.4) | 3.1 ± 0.2 | -1.1 | 10.2 ± 0.39 | 0.26 | 2.1 ± 0.2 (5.8 ± 0.6) | 2.8 ± 0.1 | -1.4 | 9.42 ± 0.05 | -0.14 |
| 6.55 | H349A | 3.8 ± 0.2 (10 ± 0.1) | 2.7 ± 0.1 | -1.5 | 8.93 ± 0.4 | -1.01 | N.D. | N.D. | N.D. | N.D. | N.D. |
| 7.35 | Y365A | 2.4 ± 0.3 (7.4 ± 1) | 3 ± 0.1 | -1.2 | 9.57 ± 0.61 | -0.37 | 2.2 ± 0.2 (6.1 ± 0.8) | 2.7 ± 0.1 | -1.5 | 9.36 ± 0.1 | -0.20 |
| 7.38 | T368A | 1.9 ± 0.2 (6.7 ± 0.9) | 3.4 ± 0.1 | -0.8 | 10.29 ± 0.42 | 0.35 | 1.7 ± 0.1 (5.2 ± 0.8) | 3.0 ± 0.2 | -1.2 | 9.73 ± 0.03 | 0.17 |
| 7.39 | T369A | 2.7 ± 0.4 (8.1 ± 1.5) | 3 ± 0.1 | -1.2 | 10.15 ± 0.41 | 0.21 | 2.5 ± 0.3 (7.4 ± 1.5) | 2.9 ± 0.2 | -1.3 | 9.83 ± 0.09 | 0.27 |
| 7.43 | Y373A | 3.1 ± 0.1 (4.4 ± 0.1) | 1.4 ± 0 | -2.8 | 5.85 ± 0.13 | -4.09 | N.D. | N.D. | N.D. | N.D. | N.D. |

**Table S5.**
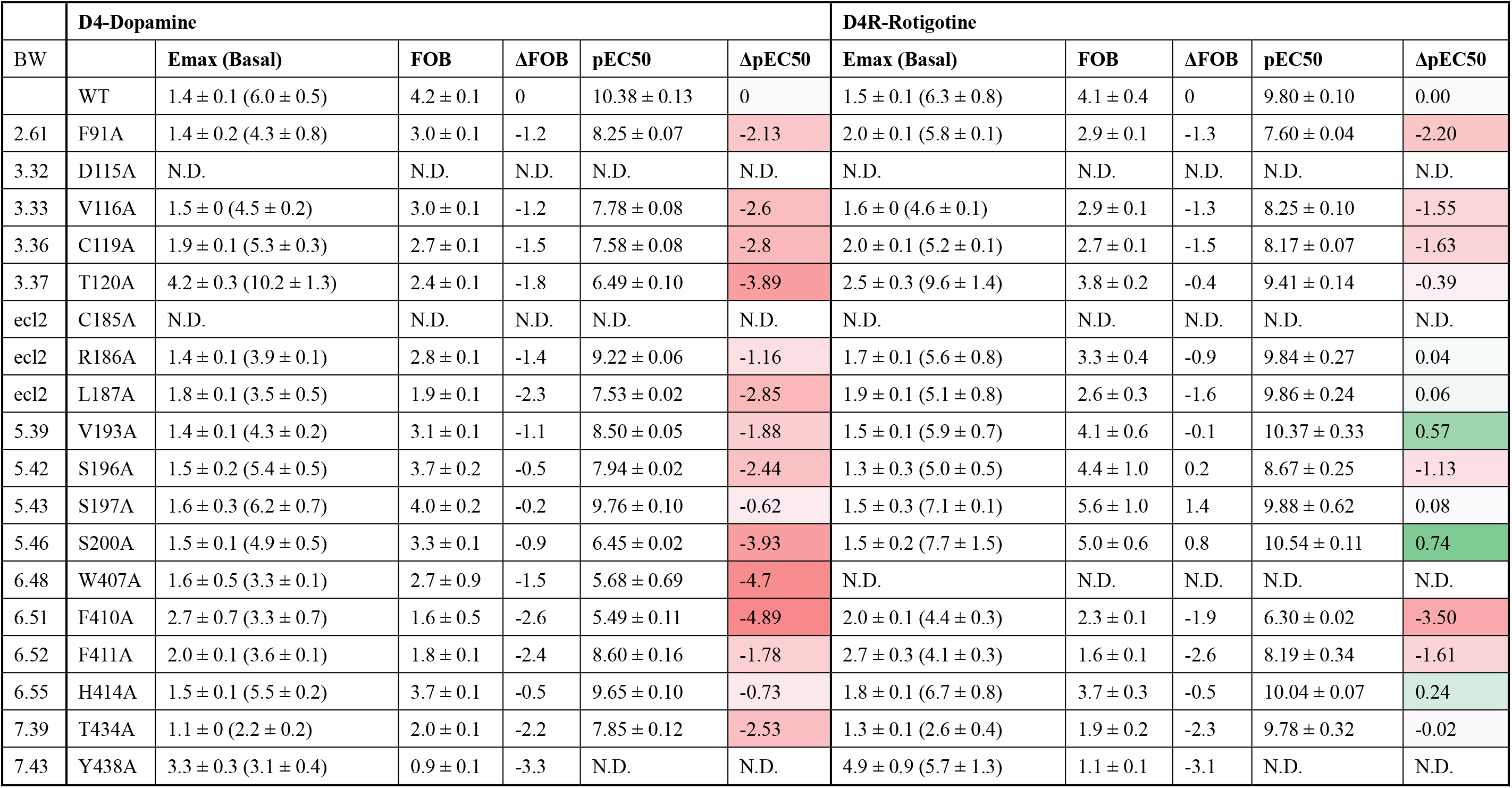
Gi-mediated cAMP accumulation assay results of D4R. Potency (pEC50) were extracted from a minimum of 3 independent assays in at least triplicate. pEC50 displayed values are mean ± SEM. Delta FOB for difference in either Fold of Basal (FOB) or Delta pEC50 when compared to wild-type receptor value. Average Emax and basal values were determined from “log(agonist) vs. response – Variable slope (four parameters) or log(inhibition) vs. response – Variable slope (four parameters)” function in Graphpad Prism 8.4 software (Graphpad Software Inc., San Diego, CA) and were divided by 10^3^ for display, basal values are enclosed with parentheses in the table column. Color scheme is based on the effects of mutations on relative pEC50 and Fold of Basal (FOB) values with red for reduced potency/efficacy and blue for increased potency/efficacy when compared to wild-type values for each ligand. ND - not determined.

**Table S6.**
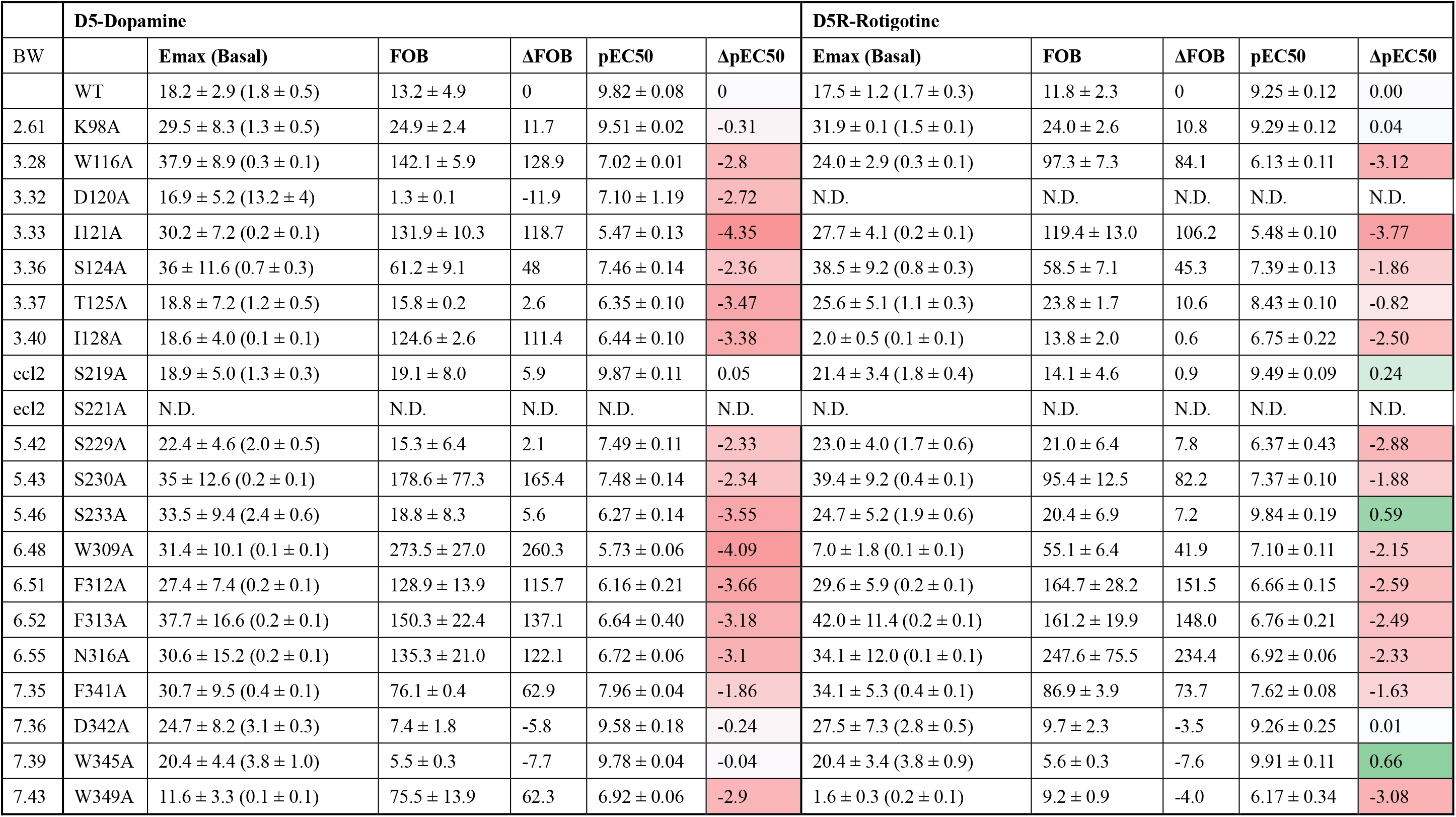
Gs-mediated cAMP accumulation assay results of D5R. Potency (pEC50) were extracted from a minimum of 3 independent assays in at least triplicate. pEC50 displayed values are mean ± SEM. Delta FOB for difference in either Fold of Basal (FOB) or Delta pEC50 when compared to wild-type receptor value. Average Emax and basal values were determined from “log(agonist) vs. response – Variable slope (four parameters) or log(inhibition) vs. response – Variable slope (four parameters)” function in Graphpad Prism 8.4 software (Graphpad Software Inc., San Diego, CA) and were divided by 10^3^ for display, basal values are enclosed with parentheses in the table column. Color scheme is based on the effects of mutations on relative pEC50 and Fold of Basal (FOB) values with red for reduced potency/efficacy and blue for increased potency/efficacy when compared to wild-type values for each ligand. ND - not determined.

**Table S8.** The distance of W^6.48^ (Toggle switch) to I^3.40^ (PIF motif) in structures of dopamine receptors.

| Receptor | Ligand |  | Signal protein | Method | PDB | Res. | State | Degree active (%) | W6.48-I3.40 distance | Reference |
| --- | --- | --- | --- | --- | --- | --- | --- | --- | --- | --- |
|  | Name | Function |  |  |  |  |  |  |  |  |
| DRD1 | Rotigotine | Agonist | Gs | cryo-EM |  | 3.2 | Active | - | 3.9 | This paper |
| DRD2 | Rotigotine | Agonist | Gi/o | cryo-EM |  | 3.0 | Active | - | 4.0 | This paper |
| DRD3 | Rotigotine | Agonist | Gi/o | cryo-EM |  | 2.7 | Active | - | 4.0 | This paper |
| DRD4 | Rotigotine | Agonist | Gi/o | cryo-EM |  | 3.2 | Active | - | 4.0 | This paper |
| DRD5 | Rotigotine | Agonist | Gs | cryo-EM |  | 3.1 | Active | - | 4.0 | This paper |
| DRD1 | Cpd1 | Agonist | Gs | X-ray | 7JOZ | 3.8 | Active | 100 | 4.2 | - |
| DRD1 | PW0464 | Agonist | Gs | cryo-EM | 7CKY | 3.2 | Active | 100 | 3.6 | 10.1016/J.CELL.2021.01.028 |
| DRD1 | dopamine | Agonist | Gs | cryo-EM | 7LJD | 3.2 | Active | 100 | 4.1 | 10.1101/2021.02.07.430101 |
| DRD1 | SKF-82526 | Agonist | Gs | cryo-EM | 7CKW | 3.2 | Active | 100 | 3.5 | 10.1016/J.CELL.2021.01.028 |
| DRD1 | CHEMBL1416789 | Agonist | Gs | cryo-EM | 7CRH | 3.3 | Active | 100 | 3.7 | 10.1016/J.CELL.2021.01.028 |
| DRD1 | dopamine | Agonist | Gs | cryo-EM | 7CKZ | 3.1 | Active | 100 | 3.5 | 10.1016/J.CELL.2021.01.028 |
| DRD1 | AA77636 | Agonist | Gs | cryo-EM | 7CKX | 3.5 | Active | 100 | 3.6 | 10.1016/J.CELL.2021.01.028 |
| DRD1 | SKF-81297 | Agonist | Gs | cryo-EM | 7LJC | 3.0 | Active | 100 | 3.8 | 10.1101/2021.02.07.430101 |
| DRD1 | 39YC3L0ZU | Agonist | Gs | cryo-EM | 7JVP | 2.9 | Active | 100 | 3.9 | 10.1016/J.CELL.2021.01.027 |
| DRD1 | apomorphine | Agonist | Gs | cryo-EM | 7JVQ | 3.0 | Active | 100 | 4.2 | 10.1016/J.CELL.2021.01.027 |
| DRD1 | SKF-81297 | Agonist | Gs | cryo-EM | 7JV5 | 3.0 | Active | 100 | 3.5 | 10.1016/J.CELL.2021.01.027 |
| DRD2 | bromocriptine | Agonist | Gi/o | cryo-EM | 7JVR | 2.8 | Active | 100 | 4.0 | 10.1016/J.CELL.2021.01.027 |
| DRD2 | bromocriptine | Agonist | Gi/o | cryo-EM | 6VMS | 3.8 | Active | 100 | 4.6 | 10.1038/S41586-020-2379-5 |
| DRD3 | PD128907 | Agonist | Gi/o | cryo-EM | 7CMV | 2.7 | Active | 100 | 4.1 | 10.1016/J.MOLCEL.2021.01.003 |
| DRD3 | pramipexole | Agonist | Gi/o | cryo-EM | 7CMU | 3.0 | Active | 100 | 3.8 | 10.1016/J.MOLCEL.2021.01.003 |
| DRD2 | spiperone | Antagonist | - | X-ray | 7DFP | 3.1 | Inactive | 1 | 8.4 | 10.1038/S41467-020-20221-0 |
| DRD2 | haloperidol | Antagonist | - | X-ray | 6LUQ | 3.1 | Inactive | 1 | 8.3 | 10.1038/S41467-020-14884-Y |
| DRD2 | rispoeridone | Inverse agonist | - | X-ray | 6CM4 | 2.9 | Inactive | 1 | 8.6 | 10.1038/NATURE25758 |
| DRD3 | eticlopride | Antagonist | - | X-ray | 3PBL | 2.9 | Inactive | 1 | 8.0 | 10.1126/SCIENCE.1197410 |
| DRD4 | L745870 | Antagonist | - | X-ray | 6IQL | 3.5 | Inactive | 83 | 4.9 | 10.7554/ELIFE.48822 |
| DRD4 | [3H]NEMONAPRIDE | Antagonist | - | X-ray | 5WIU | 2.0 | Inactive | 1 | 6.0 | 10.1126/SCIENCE.AAN5468 |
| DRD4 | [3H]NEMONAPRIDE | Antagonist | - | X-ray | 5WIV | 2.1 | Inactive | 1 | 6.1 | 10.1126/SCIENCE.AAN5468 |

**Table S7.**
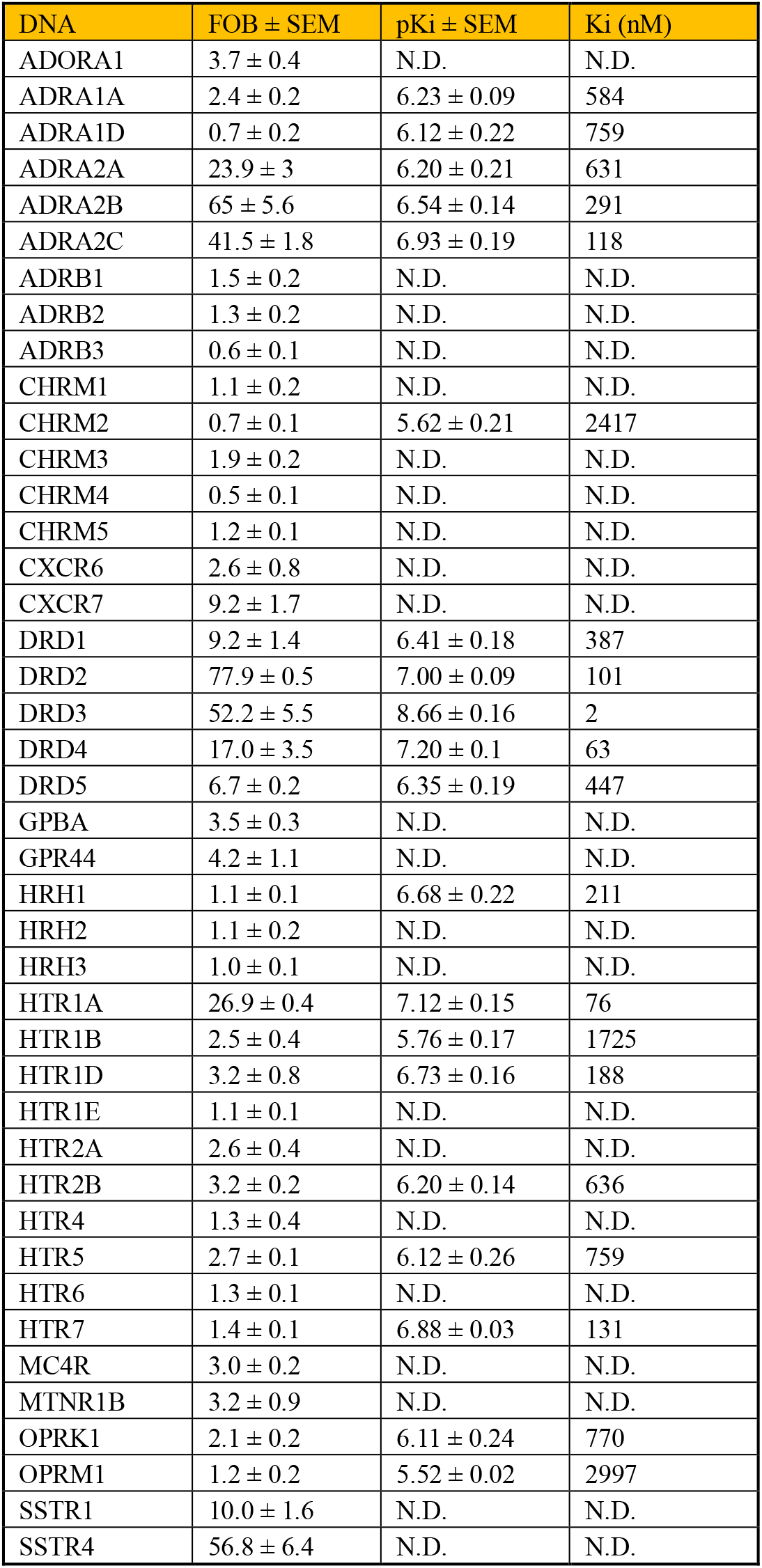
GPCRome and Radioligand Binding Assay Results. Select results of GPCRome screening and competition binding experiments of the selected receptors. Fold of Basal (FOB) and pKi values were extracted from a minimum of 3 independent assays in at least triplicate Fold of Basal (FOB) and pKi displayed values are mean ± SEM. N.D. - not determined.

| DNA | FOB ± SEM | pKi ± SEM | Ki (nM) |
| --- | --- | --- | --- |
| ADORA1 | 3.7 ± 0.4 | N.D. | N.D. |
| ADRA1A | 2.4 ± 0.2 | 6.23 ± 0.09 | 584 |
| ADRA1D | 0.7 ± 0.2 | 6.12 ± 0.22 | 759 |
| ADRA2A | 23.9 ± 3 | 6.20 ± 0.21 | 631 |
| ADRA2B | 65 ± 5.6 | 6.54 ± 0.14 | 291 |
| ADRA2C | 41.5 ± 1.8 | 6.93 ± 0.19 | 118 |
| ADRB1 | 1.5 ± 0.2 | N.D. | N.D. |
| ADRB2 | 1.3 ± 0.2 | N.D. | N.D. |
| ADRB3 | 0.6 ± 0.1 | N.D. | N.D. |
| CHRM1 | 1.1 ± 0.2 | N.D. | N.D. |
| CHRM2 | 0.7 ± 0.1 | 5.62 ± 0.21 | 2417 |
| CHRM3 | 1.9 ± 0.2 | N.D. | N.D. |
| CHRM4 | 0.5 ± 0.1 | N.D. | N.D. |
| CHRM5 | 1.2 ± 0.1 | N.D. | N.D. |
| CXCR6 | 2.6 ± 0.8 | N.D. | N.D. |
| CXCR7 | 9.2 ± 1.7 | N.D. | N.D. |
| DRD1 | 9.2 ± 1.4 | 6.41 ± 0.18 | 387 |
| DRD2 | 77.9 ± 0.5 | 7.00 ± 0.09 | 101 |
| DRD3 | 52.2 ± 5.5 | 8.66 ± 0.16 | 2 |
| DRD4 | 17.0 ± 3.5 | 7.20 ± 0.1 | 63 |
| DRD5 | 6.7 ± 0.2 | 6.35 ± 0.19 | 447 |
| GPBA | 3.5 ± 0.3 | N.D. | N.D. |
| GPR44 | 4.2 ± 1.1 | N.D. | N.D. |
| HRH1 | 1.1 ± 0.1 | 6.68 ± 0.22 | 211 |
| HRH2 | 1.1 ± 0.2 | N.D. | N.D. |
| HRH3 | 1.0 ± 0.1 | N.D. | N.D. |
| HTR1A | 26.9 ± 0.4 | 7.12 ± 0.15 | 76 |
| HTR1B | 2.5 ± 0.4 | 5.76 ± 0.17 | 1725 |
| HTR1D | 3.2 ± 0.8 | 6.73 ± 0.16 | 188 |
| HTR1E | 1.1 ± 0.1 | N.D. | N.D. |
| HTR2A | 2.6 ± 0.4 | N.D. | N.D. |
| HTR2B | 3.2 ± 0.2 | 6.20 ± 0.14 | 636 |
| HTR4 | 1.3 ± 0.4 | N.D. | N.D. |
| HTR5 | 2.7 ± 0.1 | 6.12 ± 0.26 | 759 |
| HTR6 | 1.3 ± 0.1 | N.D. | N.D. |
| HTR7 | 1.4 ± 0.1 | 6.88 ± 0.03 | 131 |
| MC4R | 3.0 ± 0.2 | N.D. | N.D. |
| MTNR1B | 3.2 ± 0.9 | N.D. | N.D. |
| OPRK1 | 2.1 ± 0.2 | 6.11 ± 0.24 | 770 |
| OPRM1 | 1.2 ± 0.2 | 5.52 ± 0.02 | 2997 |
| SSTR1 | 10.0 ± 1.6 | N.D. | N.D. |
| SSTR4 | 56.8 ± 6.4 | N.D. | N.D. |

## Notes

### Competing Interest Statement

The authors have declared no competing interest.

